# Cortical microtubules oppose actomyosin-driven membrane ingression during *C. elegans* meiosis I polar body extrusion

**DOI:** 10.1101/2023.05.26.542508

**Authors:** Alyssa R. Quiogue, Eisuke Sumiyoshi, Adam Fries, Chien-Hui Chuang, Bruce Bowerman

## Abstract

During *C. elegans* oocyte meiosis I, cortical actomyosin is locally remodeled to assemble a contractile ring near the spindle. In contrast to mitosis, when most cortical actomyosin converges into a contractile ring, the small oocyte ring forms within and remains part of a much larger and actively contractile cortical actomyosin network. This network both mediates contractile ring dynamics and generates shallow ingressions throughout the oocyte cortex during polar body extrusion. Based on our analysis of requirements for CLS-2, a member of the CLASP family of proteins that stabilize microtubules, we recently proposed that a balance of actomyosin-mediated tension and microtubule-mediated stiffness are required for contractile ring assembly within the oocyte cortical actomyosin network. Here, using live cell imaging and fluorescent protein fusions, we show that CLS-2 is part of a complex of kinetochore proteins, including the scaffold KNL-1 and the kinase BUB-1, that also co-localize to patches distributed throughout the oocyte cortex during meiosis I. By reducing their function, we further show that KNL-1 and BUB-1, like CLS-2, are required for cortical microtubule stability, to limit membrane ingression throughout the oocyte, and for meiotic contractile ring assembly and polar body extrusion. Moreover, nocodazole or taxol treatment to destabilize or stabilize oocyte microtubules, respectively, leads to excess or decreased membrane ingression throughout the oocyte and defective polar body extrusion. Finally, genetic backgrounds that elevate cortical microtubule levels suppress the excess membrane ingression in *cls-2* mutant oocytes. These results support our hypothesis that CLS-2, as part of a sub-complex of kinetochore proteins that also co-localize to patches throughout the oocyte cortex, stabilizes microtubules to stiffen the oocyte cortex and limit membrane ingression throughout the oocyte, thereby facilitating contractile ring dynamics and the successful completion of polar body extrusion during meiosis I.

## Introduction

Animal cell shape and morphogenesis are influenced by both tension and elasticity within the cell cortex (1–3). Cortical tension (contractile force) and elasticity (stiffness) both depend on the actomyosin cytoskeleton and its associated proteins. Non-muscle myosin and the architecture of cortical actin microfilaments are largely responsible for generating tension (4), while the cross-linking of cortical microfilaments to the plasma membrane by Ezrin/Radixin/Moesin (ERM) proteins generates decreased elasticity and hence increased cortical stiffness during the cell rounding associated with mitosis (5–7). Cortical microtubules appear to control cell shape and morphogenesis during plant development (8–10), and there is growing evidence for crosstalk between the microtubule and microfilament cytoskeletons (11,12). However, microtubules are generally viewed as constituting a distinct cytoskeleton that is not part of the animal cell cortex.

In contrast to the prevailing view that microtubules only interact with but are not part of animal cell cortices (1,3,11), we recently proposed that microtubules stiffen the *C. elegans* oocyte cortex to limit actomyosin-driven membrane ingression throughout the oocyte surface during meiotic cytokinesis and the extrusion of discarded chromosomes into a small polar body (13). This hypothesis followed from investigating the requirements during oocyte meiosis I cell division for CLS-2, a *C. elegans* member of the widely conserved CLASP family of proteins that bind to microtubule plus ends, promoting stability through TOG domains that bind tubulins (14–16).

During oocyte meiotic cell division, CLS-2 localizes to spindle microtubules, to kinetochores, and to small patches, also called linear elements or rods, distributed throughout the assembling spindle and the oocyte cortex (Monen et al. 2005; Dumont, Oegema, and Desai 2010; Bel Borja et al. 2020). We previously reported two prominent defects during meiosis I cell division in mutant oocytes produced by worms homozygous for null mutations in the *cls-2* locus (13). First, spindle microtubule levels were reduced, spindles never become bipolar, and chromosomes failed to separate or be extruded into a polar body. Second, in addition to the spindle defects, abnormally deep membrane ingressions occurred throughout the entire oocyte, even though the cortical actomyosin network, which extends throughout the oocyte, appeared normal. Thus CLS-2 has at least two spatially distinct functions during oocyte meiosis cell division: (i) promoting spindle microtubule stability and bipolar spindle assembly, and (ii) limiting the extent of membrane ingression throughout the oocyte during polar body extrusion.

While the dynamics of, and the requirements for, cortical actomyosin during meiotic cell division and polar body extrusion in *C. elegans* have been investigated more extensively (21–25), microtubules also are present throughout the oocyte cortex during meiosis I (13,18,26–28). However, the functional requirements for oocyte cortical microtubules in *C. elegans* have received little attention. Because CLS-2 orthologs are known to promote microtubule stability, and no differences in cortical actomyosin dynamics were detected in *cls-2* mutants compared to control oocytes (13), we hypothesized that the cortical CLS-2 patches act to stabilize microtubules and thereby directly stiffen the cortex to oppose membrane ingression. Global regulation of tension and elasticity throughout the oocyte cortex may be important for proper assembly and ingression of the small, spindle-associated contractile ring, which forms as part of the much larger cortical actomyosin network (Fig 1; see Discussion).

**Figure 1.**
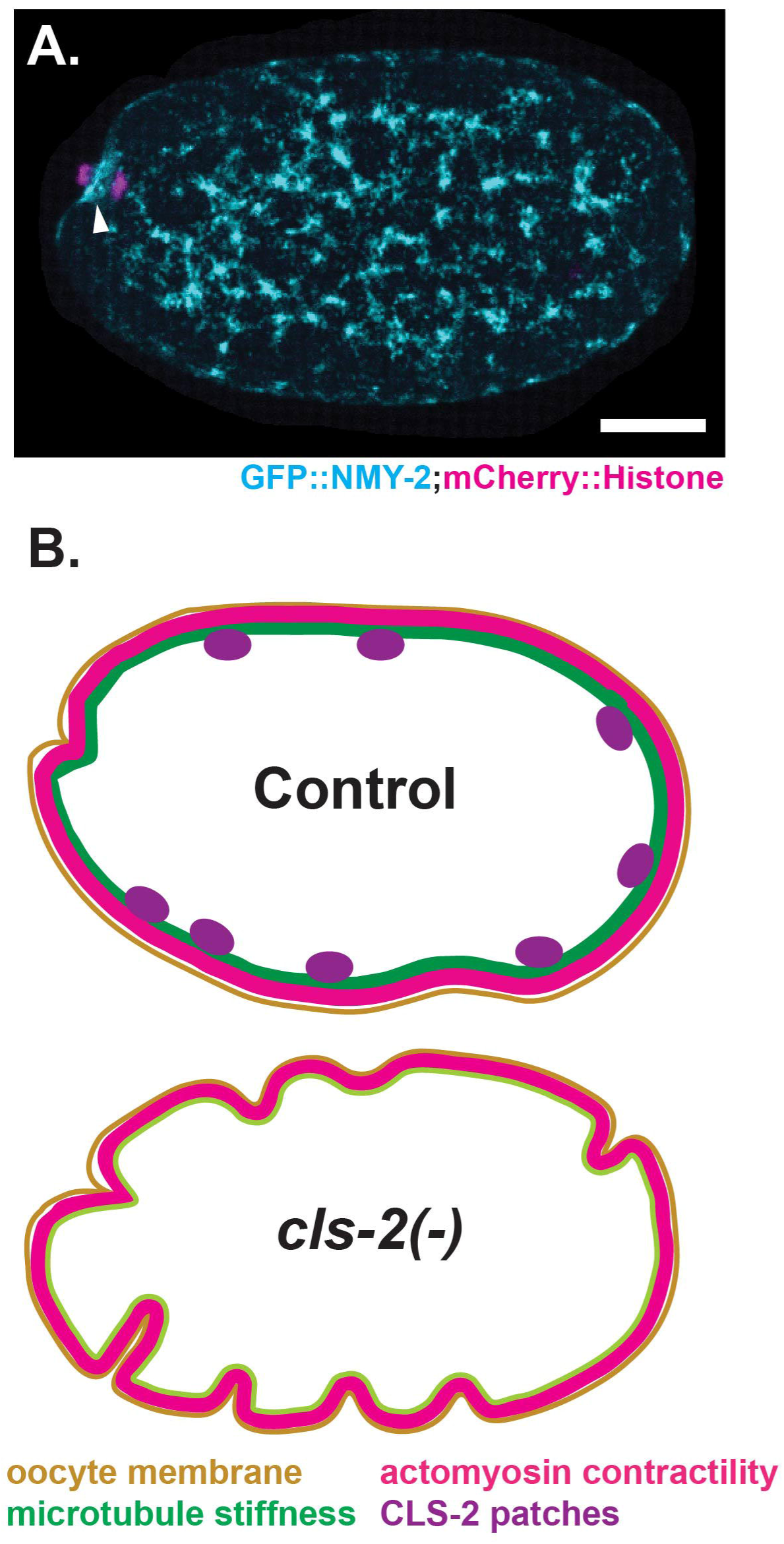
Model in which *C. elegans* polar body extrusion depends on microtubules that stiffen the cortex to limit actomyosin-driven membrane ingression throughout the oocyte. (A) Merged maximum intensity projection during meiosis I anaphase of a live *ex utero* oocyte expressing NMY-2::GFP (cyan) and mCherry::H2B (magenta) to mark non-muscle myosin and chromosomes, respectively (see Materials and Methods). Maximum intensity projection of four cortical focal planes showing non-muscle myosin (cyan) are merged with maximum intensity projections of four consecutive focal planes that encompass most of the oocyte chromosomes (magenta), which are visible at the left, anterior end of the oocyte. The oocyte cortex comprises an extensive and dynamic actomyosin network. During polar body extrusion, the network includes a contractile ring (arrowhead) that assembles near the oocyte chromosomes and ultimately constricts between separating chromosomes. (B, C) Model with schematics that depict (B) cortical CLS-2 patches (purple) stabilizing cortical microtubules to promote cortical stiffness (green) that limits membrane ingressions (tan) throughout a control oocyte by opposing cortical actomyosin-driven contractility (red); and (C) a null *cls-2* mutant oocyte with increased membrane ingression due to reduced cortical stiffness caused by reduced cortical microtubule stability, with an unchanged level of cortical actomyosin contractility. We hypothesize that a balance of stiffness and tension throughout the oocyte cortex is required for proper assembly and dynamics of the small meiotic contractile ring within the much larger cortical actomyosin network. Scale bar = 10 µm.

Other observations also suggest that microtubules are a functional component of the *C. elegans* oocyte cortex. In addition to CLS-2, two other widely conserved regulators of microtubule stability are known to influence oocyte cortical microtubule levels: the kinesin 13/MCAK microtubule depolymerase family member called KLP-7, and another TOG domain protein, ZYG-9/chTOG. Both KLP-7 and ZYG-9 are required for oocyte meiotic spindle assembly, and reducing the function of either results in elevated levels of both spindle and cortical microtubules (27–29). While the consequences of increased cortical microtubules in *klp-7* and *zyg-9* mutant oocytes have not been investigated, microtubules clearly are a regulated component of the oocyte cortex and therefore might influence cortical elasticity and stiffness, and contractile ring dynamics, during meiotic cell division.

In addition to its roles in oocyte meiotic spindle assembly and polar body extrusion, CLS-2 also acts at kinetochores early in mitosis to stabilize microtubule/kinetochore attachments (30–32). In both meiotic oocytes and mitotic blastomeres, CLS-2 is part of an outer kinetochore sub-complex that includes the spindle assembly checkpoint kinase BUB-1 and the redundant coiled-coil proteins and CENP-F orthologs HCP-1 and -2.

Like other outer kinetochore sub-complexes, the BUB-1/HCP-1/2/CLS-2 sub-complex depends on the kinetochore scaffolding protein KNL-1 for its kinetochore localization (19,30). Intriguingly, like CLS-2, both KNL-1 and BUB-1 also localize to linear elements that are enriched within the assembling spindle and are distributed throughout the oocyte cortex during meiosis I cell division (18–20). While all three proteins co-localize to kinetochores, it is not known whether they also co-localize within the patches, or linear elements, distributed throughout the oocyte cortex.

The spindle-associated linear elements are thought to promote kinetochore expansion and hence the capture of spindle microtubules during mitotic cell division, and similar structures have been described in mammalian cells (33). The role(s) played by the cortical oocyte patches are not known, although KNL-1 linear elements were shown not to co-localize with cortical microtubules in fixed oocytes (18). Here we report our investigation of requirements for KNL-1, BUB-1, CLS-2, and cortical microtubules during *C. elegans* polar body extrusion. Our results support the hypothesis that microtubules stiffen the oocyte cortex and thereby limit membrane ingression during *C. elegans* meiosis I polar body extrusion.

## Results

### The kinetochore proteins KNL-1, BUB-1 and CLS-2 co-localize to linear elements, or patches, distributed throughout the oocyte cortex during meiosis I

To determine when KNL-1, BUB-1 and CLS-2 are present, and if they co-localize to cortical patches, we used spinning disk confocal microscopy and live cell imaging to track fluorescent protein fusions to each protein (see Materials and Methods). First, we examined their localization throughout meiosis I and II, after imaging *in utero* oocytes that express a GFP fusion to one of the three kinetochore proteins, and mCherry fused to a histone (mCherry::H2B) to mark chromosomes. As reported previously (Monen et al 2005; Dumont et al 2010; Schlientz 2020), we detected the GFP fusions to KNL-1, BUB-1 and CLS-2 shortly after nuclear envelope breakdown during meiosis I, associated with chromosomes and also in linear elements scattered throughout the assembling spindle and less densely throughout the oocyte cortex (Figs 2, S1, Movie 1). The KNL-1, BUB-1, and CLS-2 cortical structures persisted until the beginning of anaphase B, when they became undetectable, approximately 13 minutes after nuclear envelope breakdown.

**Figure 2.**
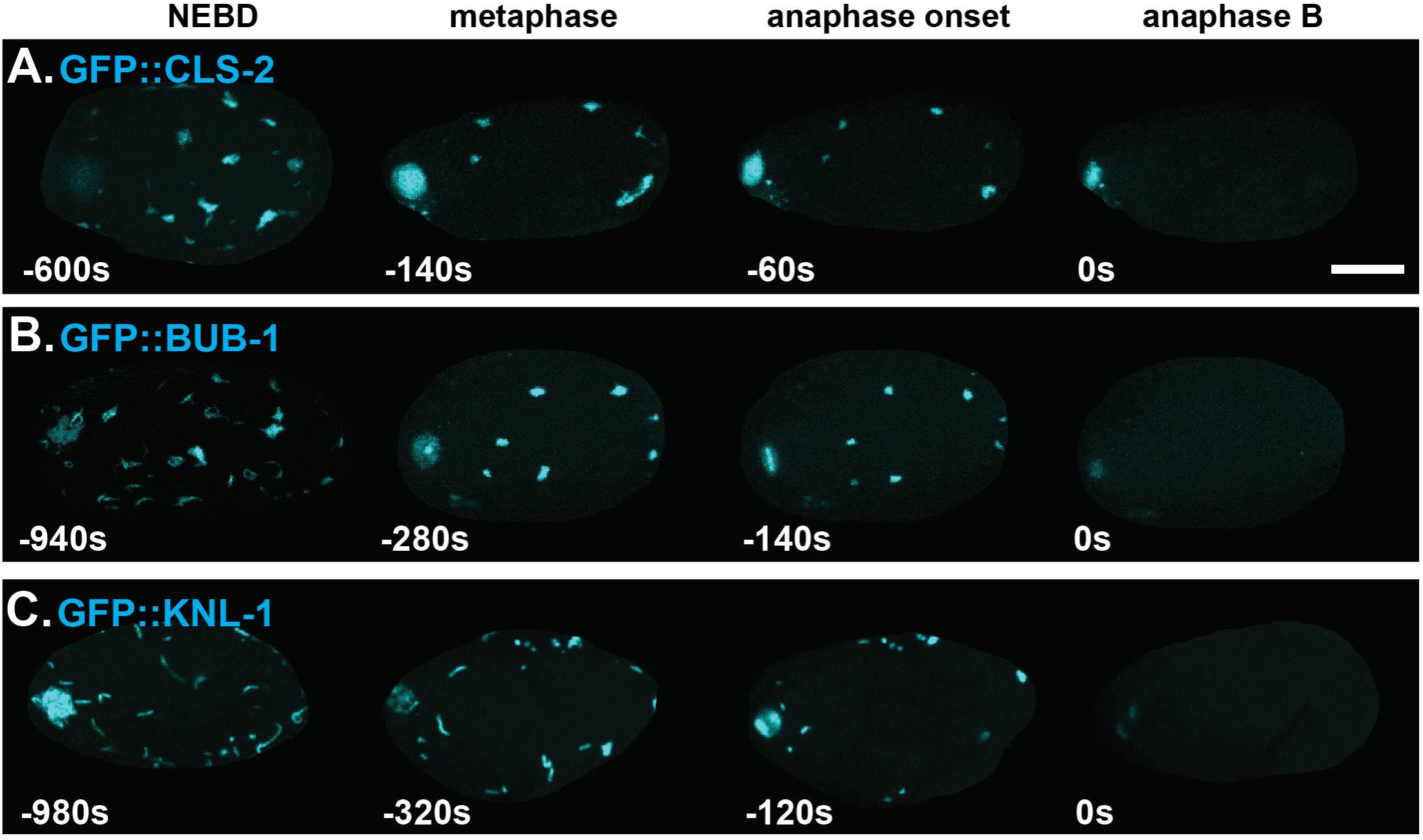
CLS-1, BUB-1 and KNL-1 are present in patches distributed throughout the oocyte cortex during meiosis I. Maximum intensity projections of four cortical focal planes during meiosis I in live *ex utero* oocytes expressing mCherry::H2B to mark chromosomes (not shown) and GFP fusions to CLS-2 (A), BUB-1 (B), and KNL-1 (C) in cyan. All three kinetochore proteins were present in linear elements associated with the oocyte spindle and chromosomes (visible at left, anterior end in most oocytes) and also distributed throughout the cortex, beginning at nuclear envelope breakdown (NEBD). The cortical linear elements became more patch-like over time and persisted until the beginning of anaphase B, when they were no longer detectable. t = 0 seconds denotes the beginning anaphase B, when chromosomes were most condensed (see Materials and Methods). Scale bar = 10 µm.

These cortical structures varied in size and shape, were mobile, sometimes fusing together or fragmenting into pieces both near the spindle and at the cortex (Movie 1), and they generally became less linear and more patch-like as meiosis I proceeded (Figs 2, S1, Movie 1). Hereafter, we refer to these cortical structures as patches. As documented previously (Monen et al 2005, Dumont et al 2010), as the KNL-1, BUB-1 and CLS-2 cortical patches dissipated during anaphase, BUB-1 and CLS-2 but not KNL-1 localized to the central spindle between the separating meiotic chromosomes, and during meiosis II, KNL-1, BUB-1 and CLS-2 all localized to chromosomes, and BUB-1 and CLS-2 to the central spindle, but no cortical patches were detected near the spindle or throughout the cortex (Figs 2, S1, Movie 1).

We next asked if KNL-1, BUB-1 and CLS-2 co-localize to cortical patches during meiosis I, in oocytes expressing mCherry::H2B to mark chromosomes and different combinations of RFP and GFP fusions to KNL-1, BUB-1, and CLS-2. Using live imaging of *ex utero* oocytes, we saw that both GFP::BUB-1 and GFP::CLS-2 co-localized with RFP::KNL-1, and RFP::BUB-1 co-localized with GFP::CLS-2, at most if not all cortical patches during meiosis I (Fig 3A, Movie 2).

**Figure 3.**
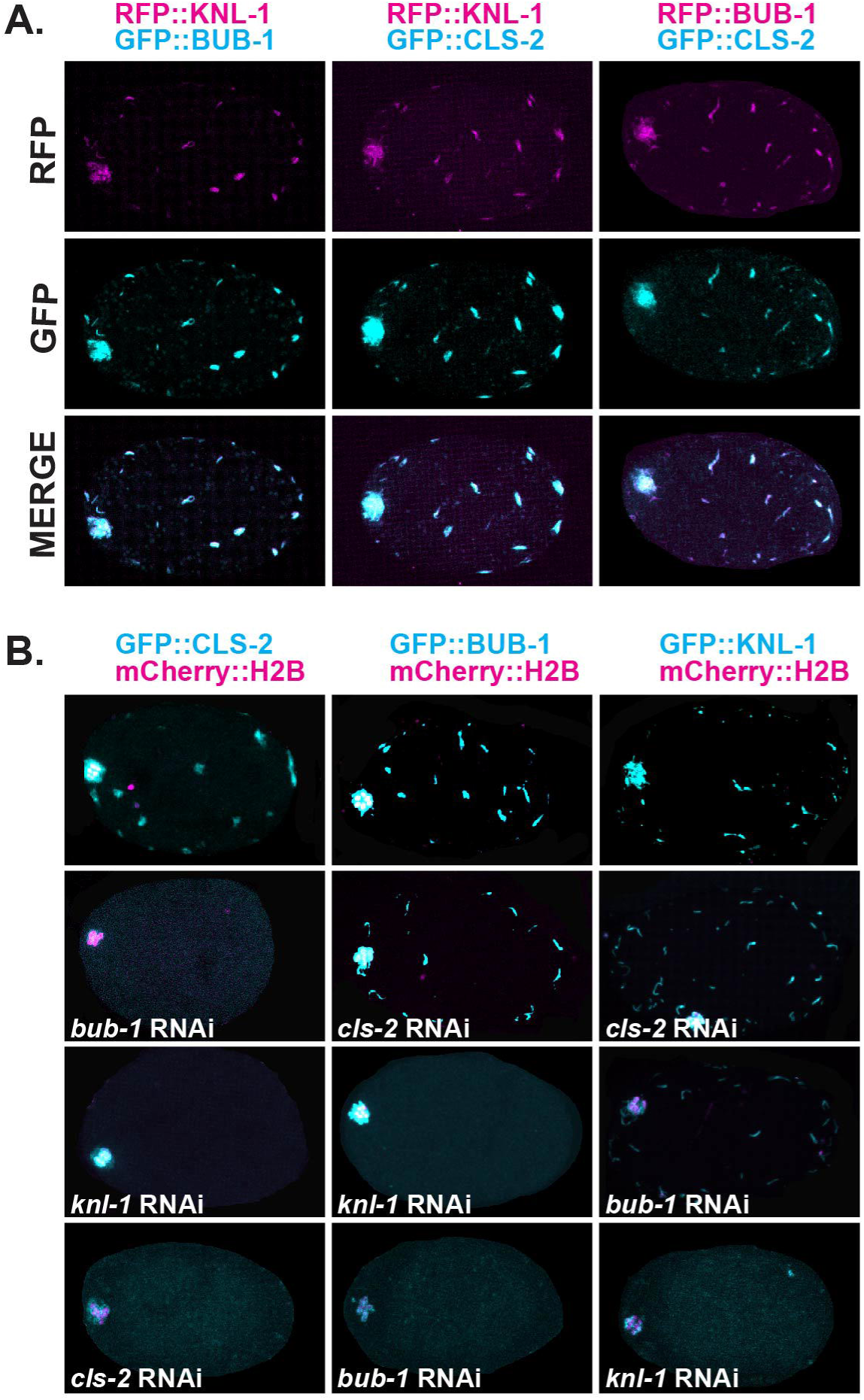
CLS-2, BUB-1 and KNL-1 exhibit the same hierarchical dependency for co-localization to cortical patches as they do to kinetochores. (A) Maximum intensity projections of 4 cortical focal planes during meiosis I in*ex utero* oocytes expressing both RFP (magenta) and GFP (cyan) fusions as follows: RFP::KNL-1 and GFP::BUB-1 (left column), RFP::KNL-1 and GFP::CLS-2 (middle column), or RFP::BUB-1 and GFP::CLS-2 (right column). All images are from oocytes at roughly metaphase in *ex utero* oocytes, when the patches are numerous and bright. Each pair of proteins co-localized to all patches. Spindle-associated GFP signal is visible at the left, anterior end of each oocyte. (B) Merged maximum intensity projections of four cortical focal planes, and four consecutive focal planes that encompass most of the oocyte chromosomes, during meiosis I in *ex utero* oocytes that all express mCherry::H2B (magenta) to mark chromosomes, and GFP (cyan) fusions to CLS-2 (left column), BUB-1 (middle column) or KNL-1 (right column). RNAi knockdowns of CLS-2, BUB-1 and KNL-1 are indicated. As at kinetochores (see text), CLS-2 required both BUB-1 and KNL-1 for localization to cortical patches, BUB-1 required only KNL-1, and KNL-1 required neither CLS-2 nor BUB-1, although there may be a partial dependence on BUB-1. As shown at bottom, our feeding RNAi protocols only partly knock down CLS-2, BUB-1 and KNL-1, with some signal remaining associated with chromosomes. Scale bars = 10 µm.

At oocyte kinetochores, CLS-2 requires both KNL-1 and BUB-1 for kinetochore localization, while BUB-1 requires only KNL-1, and KNL-1 does not depend on either BUB-1 or CLS-2 (19).To assess their dependencies at cortical patches, we used RNA interference (RNAi; see Materials and Methods) to knock down each protein in strains expressing GFP fusions to one of the other two proteins and observed the same dependency (Fig 3B). CLS-2 was undetectable at cortical patches after knocking down either KNL-1 or BUB-1, while BUB-1 depended only on KNL-1, and KNL-1 localization to cortical patches was not affected by either BUB-1 or CLS-2 knockdown. To summarize, during *C. elegans* oocyte meiosis I cell division, the kinetochore proteins KNL-1, BUB-1 and CLS-2 also co-localize to dynamic cortical patches that appear upon nuclear envelope breakdown and persist until the beginning of anaphase B during meiosis I.

### KNL-1, BUB-1 and CLS-2 are each required to limit membrane ingression during meiosis I polar body extrusion

To determine whether BUB-1 and KNL-1, like CLS-2 (13), are required to limit membrane ingression throughout the oocyte during meiosis I polar body extrusion, we examined membrane dynamics in control and in *cls-2(or1948)* null mutants, and after using RNAi to knockdown KNL-1 or BUB-1, in oocytes that express GFP fused to a plasma membrane marker (GFP::PH) and the chromosome marker mCherry::H2B (see Materials and Methods). Because chromosome separation fails in *cls-2* mutant oocytes (34) (13,16,19), we defined the beginning of anaphase B, in the absence of chromosome separation, as the timepoint when the oocyte chromosome mass was most compacted (Redderman et al. 2018). In control oocytes, spindle associated furrows began to ingress near the beginning of anaphase B and constricted as meiosis progressed. The membrane ingressions associated with polar body extrusion were accompanied by multiple shallow ingressions throughout the oocyte during anaphase B (Figs 4A, S2, Movie 3). During polar body extrusion attempts in *cls-2* mutant oocytes, abnormally deep membrane ingressions appeared throughout the oocyte, as previously reported (Schlientz and Bowerman 2020), with the extent of ingression peaking late in anaphase B and then rapidly declining (Figs 4B, S2, Movie 3). After RNAi knockdown of either KNL-1 or BUB-1, we observed abnormally deep membrane ingressions that were indistinguishable from those observed in *cls-2* mutants (Figs 4C,D, S2, Movie 3). Polar body extrusion during meiosis I failed in ∼2/3 of *cls-2(or1948)* oocytes (Fig S3B), consistent with previous reports (13). However, after either KNL-1 or BUB-1 knockdown, polar body extrusion was usually successful, with some chromosomes detected within a polar body after the completion of meiosis I in over 80% of the mutant oocytes (Fig S3B), possibly due to our feeding RNAi only partially knocking down these proteins (Fig 3B; see Materials and Methods).

**Figure 4.**
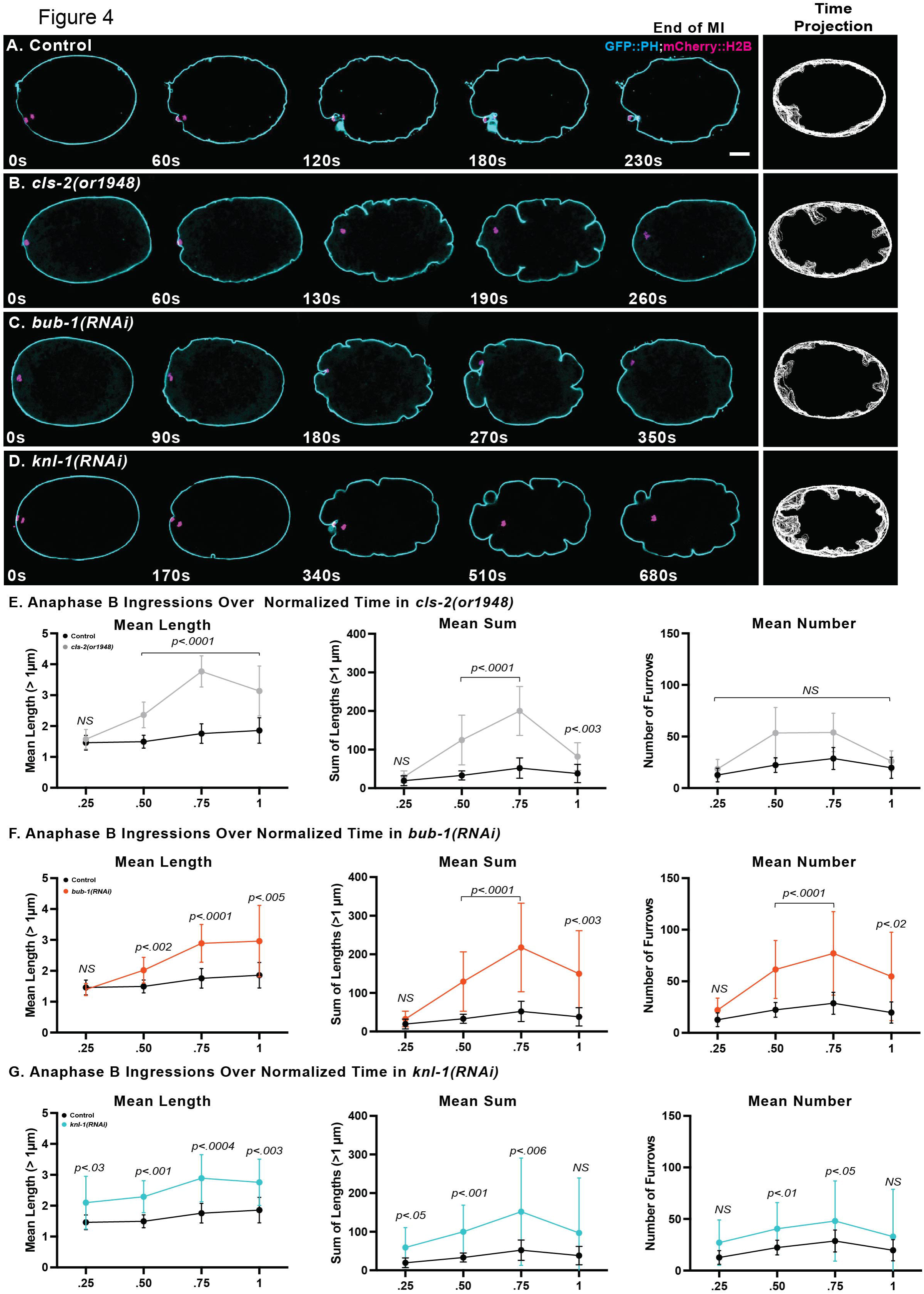
CLS-2, BUB-1, and KNL-1 each limit membrane ingression during meiosis I. (A-D) Selected and merged focal planes from *ex utero* oocytes during meiosis I anaphase B that express GFP::PH (cyan) and mCherry::H2B (magenta) to mark the plasma membrane and chromosomes, respectively. A single central focal plane is shown for the membrane, merged with a maximum intensity projection of 4 consecutive focal planes that encompass most of the oocyte chromosomes, which are visible at the left, anterior end of each oocyte. In this and all subsequent figures, unless otherwise noted, time is in seconds, with t = 0 denoting the beginning of anaphase B, when the chromosomes were most compacted, and the last time point showing the end of meiosis I, when oocyte chromosomes first began to decondense prior to meiosis II (see Materials and Methods). Time projections of all central focal plane anaphase B time points for each genotype are shown in the column at the far right (see Fig S2 for additional time projections of each genotype). Limited ingressions occurred throughout anaphase B in control oocytes (A), with excess furrowing observed in *cls-2(or1948)* null mutant oocytes (B) and after RNAi depletion of BUB-1 (C) and KNL-1 (D). Note that progression through meiosis I often took longer and was variable in *cls-2* null mutant oocytes, and after BUB-1 and KNL-1 knockdowns, relative to control oocytes (see Fig S3A for all data).(E-F) Quantification of anaphase B membrane ingressions over four equally spaced normalized anaphase B time intervals, to control for the variation in cell cycle time. Graphs depict the mean length of all membrane ingressions per oocyte that were 1μm or greater in length (left), the mean sum of all ingressions 1μm or greater in length (middle), and the mean number of ingressions 1μm or greater in length (to right), at each time point (see Materials and Methods). The mean length, and the mean sum of lengths, both peaked late in anaphase B for all genotypes. P values denote comparisons between mutant and controls at indicated time points. Note that the mean ingression length, the mean sum of lengths, and the mean ingression number were all significantly increased after both BUB-1 and KNL-1 knockdown, but only the mean length and the mean sum of lengths, not the mean number of ingressions, were significantly increased in *cls-2(or1948)* oocytes. For all figures, the Mann-Whitney U-test was used to calculate P-values (see Supplemental Files S2, S3). Error bars and values are mean ± SEM. Scale bar = 10 µm.

To quantify membrane ingression during polar body extrusion, we developed a data analysis pipeline to objectively measure the number and length of all ingressions (Figs 4E-G, S3B,C). In brief, we used a convex hull of the oocyte contour, from a single central focal plane, as a reference for measuring furrow appearance and length, and we wrote a program that quantifies the number and length of all furrows (see Materials and Methods). We manually excluded the longest furrow associated with the oocyte chromosomes, and the program excluded furrows that were less than 1 μm in length, over a normalized anaphase B timescale. The mean length of ingressions, and the mean sum of lengths, were significantly increased in *knl-1, bub-1* and *cls-2* mutants compared to control oocytes (Fig. 4E-G). We also detected a significant increase in the number of ingressions 1 μm or greater in length after both KNL-1 and BUB-1 knockdowns, but not in *cls-2(or1948)* oocytes (Fig 4E-G), even though the RNAi knockdowns only partially reduce gene function (see Fig 3B) and the *or1948* allele eliminates all *cls-2* function. This difference suggests that KNL-1/BUB-1-dependent factors other than CLS-2 contribute to the negative regulation of membrane ingression during polar body extrusion (see Discussion).

### KNL-1, BUB-1 and CLS-2 stabilize cortical microtubules during meiosis I polar body extrusion

We next asked how KNL-1, BUB-1 and CLS-2 limit the extent of membrane ingression during oocyte meiotic cell division. We previously hypothesized that the cortical CLS-2 patches stabilize cortical microtubules to promote stiffness and thereby limit actomyosin-driven membrane ingression during meiosis I polar body extrusion (Fig 1). However, cortical microtubule levels are much lower than those associated with the meiotic spindle, and we were unable in our previous study to detect a significant difference in cortical microtubule levels using relatively low resolution live imaging data from oocytes within immobilized whole mount worms (13).

To determine whether KNL-1, BUB-1 and CLS-2 are required to stabilize cortical microtubules during oocyte meiotic cell division, we have now used higher resolution *ex utero* live imaging of oocytes that express GFP fused to a β-tubulin (GFP::TBB-2) to mark microtubules, and mCherry::H2B to mark chromosomes (see Materials and Methods). In control oocytes, we observed a network of microtubule foci distributed throughout the cortex (Fig 5A, Movie 4). Consistent with our hypothesis that the KNL-1/BUB-1/CLS-2 cortical patches stabilize cortical microtubules, we observed a partial but apparent reduction of cortical microtubules in all three mutants compared to control oocytes (Figs 5B-D, Movie 4).

**Figure 5.**
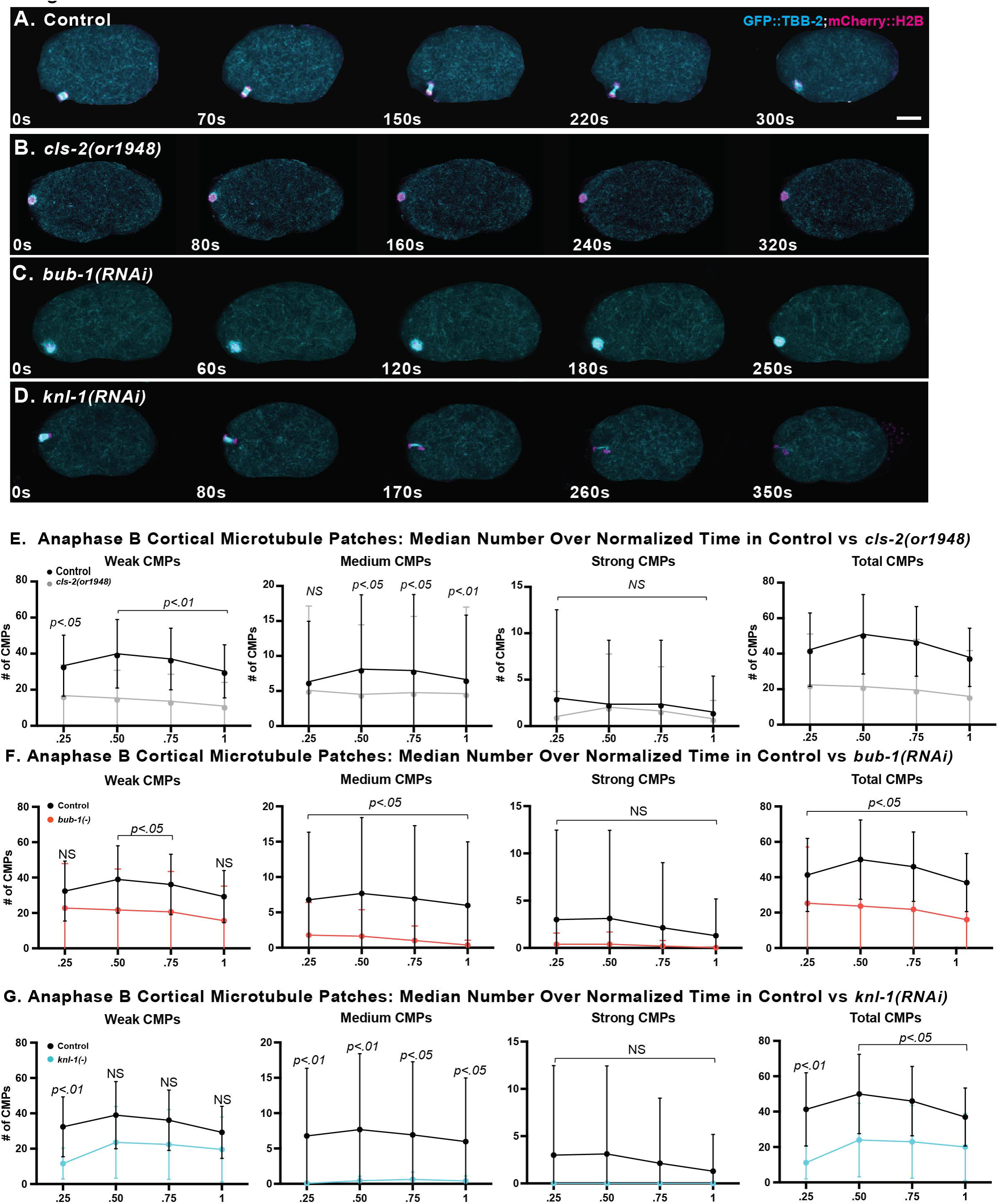
Cortical microtubules are reduced in *cls-2, bub-1,* and *knl-1* mutant oocytes. Merged maximum intensity projections during meiosis I anaphase B of *ex utero* oocytes expressing GFP::TBB-2 (cyan) and mCherry::H2B (magenta) to mark microtubules and chromosomes, respectively. Maximum intensity projections of four cortical focal planes showing microtubules are merged with maximum intensity projections of four consecutive focal planes that encompass most of the oocyte chromosomes, which are visible at the left, anterior end of each oocyte. Many small foci within an extensive network of microtubules are present in control oocytes (A), with an apparent reduction in overall signal intensity observed in all three mutants (B-D). (E-F) Quantification of mean number of weak, medium and strong cortical patches (CMPs), and total CMPs (sum of weak, medium, and strong CMPs), over normalized time in control and mutant oocytes (see Materials and Methods). The mean number of medium CMPs, and total CMPS, was significantly reduced in all three mutants. Scale bar = 10 µm.

To quantify cortical microtubules, we used Imaris software to define and count Cortical Microtubule Patches, or CMPs, during polar body extrusion (see Materials and Methods). In brief, CMPs were defined based on the local variance of pixel intensity above threshold variance values, counted throughout the cortex at each time point, and divided into weak, medium and strong CMPs based on relative pixel intensity (Fig S4, Movie 5). We observed significant decreases in the number of medium CMPs, and in the total number of all CMPs, after KNL-1 and BUB-1 RNAi knockdowns and in *cls-2(or1948)* oocytes, compared to control oocytes (Figs 5E-G). Note that while the KNL-1/BUB-1/CLS-2 cortical patches are undetectable by the beginning of anaphase B, we detected lower levels of cortical microtubules throughout anaphase B. If these cortical patches do stabilize microtubules, whether they do so prior to anaphase B, or with undetectable protein levels during anaphase B, requires further investigation (see Discussion). We conclude that KNL-1, BUB-1 and CLS-2 all promote cortical microtubule stability during meiosis I anaphase B.

### Chemical alteration of microtubule levels alters oocyte membrane ingression and interferes with polar body extrusion

If the reduced levels of oocyte cortical microtubules in *knl-1, bub-1* and *cls-2* mutant oocytes increase elasticity and thus decrease stiffness, then disruption of microtubules by treatment with nocodazole should also reduce stiffness and result in excessively deep membrane ingression throughout the oocyte during polar body extrusion. To examine the effects of nocodazole treatment, first we imaged *ex utero* oocytes that express GFP::TBB-2 and mCherry::H2B to confirm that our nocodazole treatments depleted spindle and cortical microtubules. In control oocytes exposed only to DMSO, all appeared normal but upon exposure to 10 ug/ml nocodazole (see Materials and Methods), microtubule levels were greatly reduced, both in association with oocyte chromosomes and throughout the cortex (Fig 6A, Movie 4).

**Figure 6.**
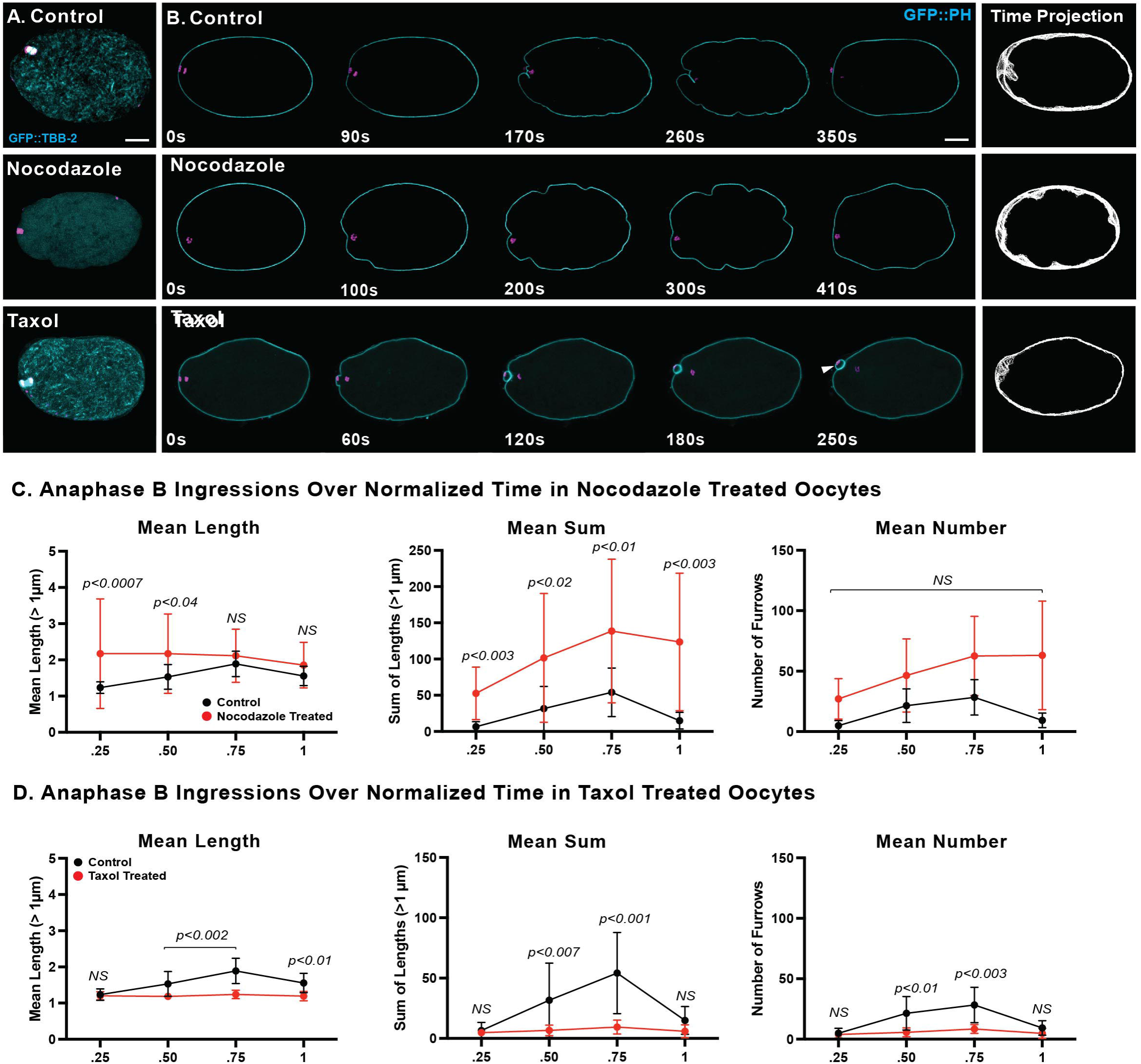
Nocodazole and taxol modulate membrane ingression during *C. elegans* oocyte meiosis I. (A) Merged maximum intensity projections during meiosis I anaphase B of *ex utero* oocytes expressing GFP::TBB-2 (cyan) and mCherry::H2B (magenta) to mark microtubules and chromosomes, respectively, in control (top), nocodazole-treated (middle), and taxol-treated (bottom) oocytes. Maximum intensity projections of four cortical focal planes showing microtubules are merged with maximum intensity projections of four consecutive focal planes that encompass most of the oocyte chromosomes, which are visible at the left, anterior end of each oocyte. (B) Selected and merged focal planes of *ex utero* oocytes expressing GFP::PH (cyan) and mCherry::H2B (magenta) to mark membranes and chromosomes, respectively, in control (top row), nocodazole-treated (middle row), and taxol-treated (bottom row) oocytes. A single central focal plane is shown for the membrane, merged with a maximum intensity projection of 4 consecutive focal planes that encompass most of the oocyte chromosomes. Time projections of single central focal planes for the membrane are shown at far right. (C-D) Quantification of the mean length of membrane ingressions that were 1μm or more in length, the mean sum of lengths, and the mean number of membrane ingressions, over normalized anaphase B time in control and chemically treated oocytes. While polar body extrusion always failed after nocodazole treatment, polar body extrusion was usually successful after taxol treatment (arrowhead; also see Fig S3B). Note that the mean length and mean sum of lengths, but not the mean number of ingressions were increased in nocodazole-treated oocytes, while both the mean length and number of ingressions were decreased in taxol-treated oocytes (see text). Scale bar = 10 µm.

To assess the impact of nocodazole-mediated microtubule depletion on oocyte membrane ingression, we imaged *ex utero* oocytes that express GFP::PH and mCherry::H2B fusions to mark the membrane and chromosomes. In control oocytes, spindle-associated furrows ingressed during chromosome separation, with shallow furrows also appearing throughout the oocyte during anaphase B. After exposure to nocodazole, ingressions still appeared near the oocyte chromosomes, but the chromosomes failed to separate, polar body extrusion failed, and abnormally deep membrane ingressions appeared throughout the oocytes (Figs 6B, S3B, Movie 3). The mean ingression length and mean sum of all lengths per oocyte were significantly increased relative to control oocytes, although ingression number was not affected (Fig 6C; see Discussion).

We next asked if chemically stabilizing oocyte microtubules with taxol would reduce the extent of membrane ingression during polar body extrusion. Taxol treatment substantially elevated cortical microtubule levels during meiosis I (Fig 6A, Movie 3), and significantly reduced the extent of membrane ingression (Figs 6B, 6D, Movie 3), although polar bodies were extruded in nearly all treated oocytes (Figs 6B, S3B). The increased and decreased membrane ingressions observed after nocodazole and taxol treatment, respectively, further support our hypothesis that cortical microtubules limit membrane ingression during meiosis I polar body extrusion.

### Increasing cortical microtubules suppresses membrane ingression in cls-2 mutants

If decreased cortical microtubule stability in *cls-2* mutant oocytes causes excess membrane ingression, then increasing cortical microtubules in *cls-2* mutants should suppress the excess ingression. Two microtubule regulators in *C. elegans,* the kinesin 13/microtubule depolymerase family member KLP-7, and the XMAP215/chTOG family member ZYG-9, both have been shown to limit spindle and cortical microtubules levels (27,28,35). We first asked whether the increased cortical microtubules in *klp-7* and *zyg-9* mutants depend on CLS-2 by quantifying cortical microtubule patches (CMP) during anaphase B in *ex utero klp-7(RNAi)* and *zyg-9(RNAi)* single mutant, and *cls-2(or1948) klp-7(RNAi)* and *zyg-9(RNAi); cls-2(or1948)* double mutant oocytes, all expressing GFP::TBB-2 and mCherry::H2B (Fig. 7A-F, Movie 4). We observed significant increases in total CMP number, relative to control oocytes, not only in *klp-7* and *zyg-9* single mutants, but also in the corresponding *cls-*2 double mutants (Fig. 7G).

**Figure 7.**
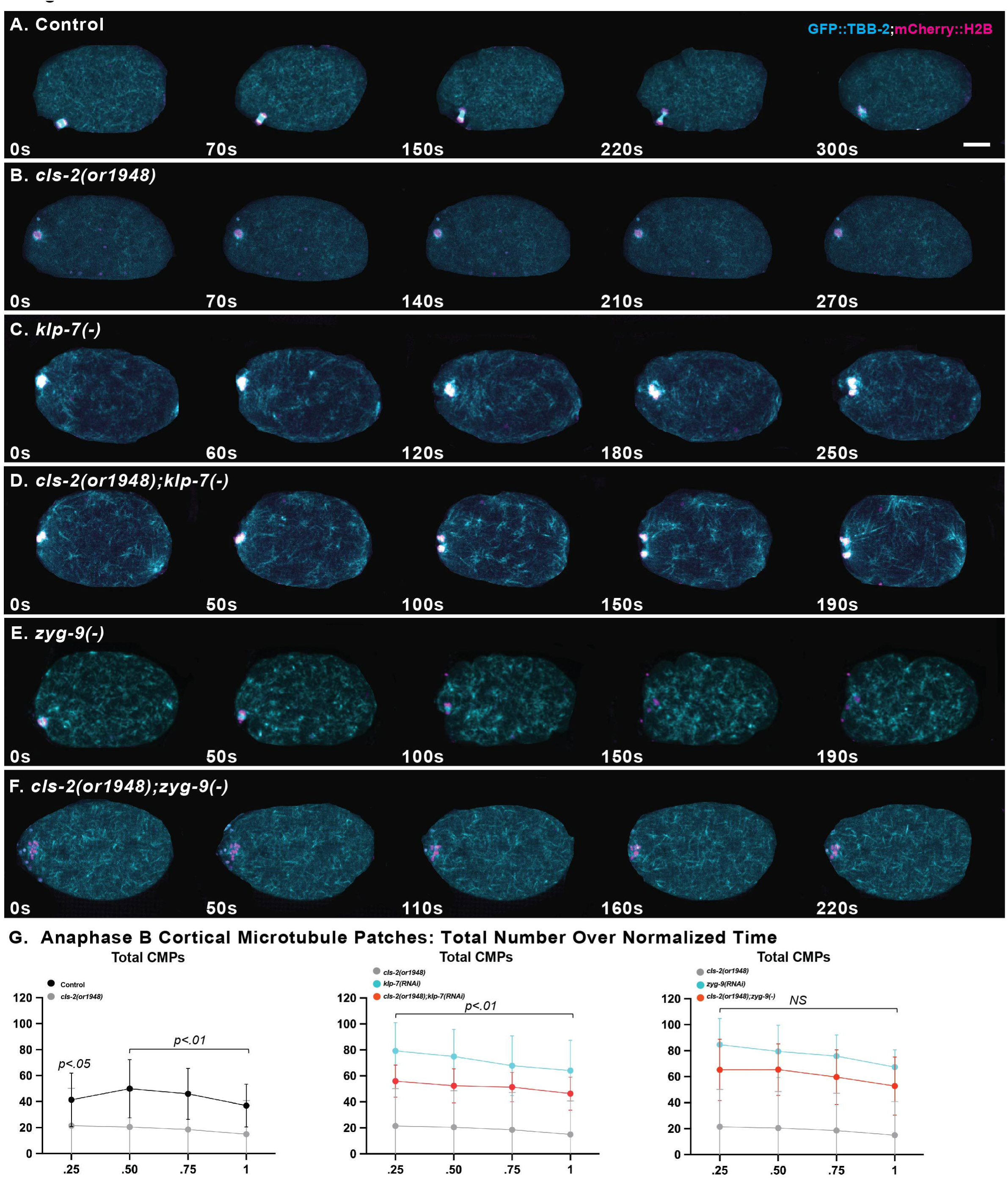
Loss of KLP-7/kinesin-13 or ZYG-9/chTOG increases cortical microtubules in*cls-2(or1948)* oocytes. (A-F) Merged maximum intensity projections during meiosis I anaphase B of live *ex utero* control and mutant oocytes expressing GFP::TBB-2 (cyan) and mCherry::H2B (magenta) to mark microtubules and chromosomes. Maximum intensity projections of four cortical focal planes showing microtubules are merged with maximum intensity projections of four consecutive focal planes that encompass most of the oocyte chromosomes. Spindle-associated microtubules and chromosomes are visible at the left, anterior end of each oocyte. (G) Total cortical microtubule patch (CMP) numbers over normalized anaphase B time. Total CMPs were significantly reduced in *cls-2(or1948)* mutants, and significantly increased in *klp-7(RNAi)* and *zyg-9(RNAi)* mutants, compared to control oocytes. Total CMPs were significantly decreased in*cls-2(or1948) klp-7* (RNAi) double mutants compared to *klp-7(RNAi)* single mutants, but not in *zyg-9(RNAi); cls-2(or1948)* double mutants compared to *zyg-9(RNAi)* single mutant oocytes. Scale bar = 10 µm.

While KLP-7 and ZYG-9 knockdowns both led to increased cortical microtubules, the microtubule properties in each mutant were distinct. The increased cortical microtubules in *klp-*7 mutants did not show a statistically significant dependence on CLS-2, but cortical microtubule levels were partially but significantly reduced in *zyg-9; cls-*2 double mutants relative to *zyg-9* single mutants. Nevertheless, cortical microtubules in both double mutants were still significantly elevated compared to control and *cls-2* mutants (Fig 7G). Finally, the distribution of cortical microtubules appeared to vary qualitatively in *klp-7* compared to *zyg-9* single mutants, and similarly in the corresponding double mutants. The cortical microtubules in *klp-7* mutant oocytes appeared more evenly distributed (Fig 7C, Movie 4), whereas the cortical microtubules in *zyg-9* mutant oocytes formed many small puncta (Fig 7E, Movie 4).

After determining that depleting KLP-7 or ZYG-9 elevates cortical microtubule levels, we next asked whether the excess membrane ingression in *cls-2* mutant oocytes is suppressed in *cls-2 klp-7* and *zyg-9; cls-2* double mutants, again imaging oocytes that express GFP::PH and mCherry::H2B (Fig 8A-F, Movie 3). We observed significant reductions in both the mean ingression length and the mean sum of lengths in *cls-2 klp-7* and *zyg-9; cls-2* double mutants compared to *cls-2* mutant oocytes (Figs 8G, 3H).

**Figure 8.**
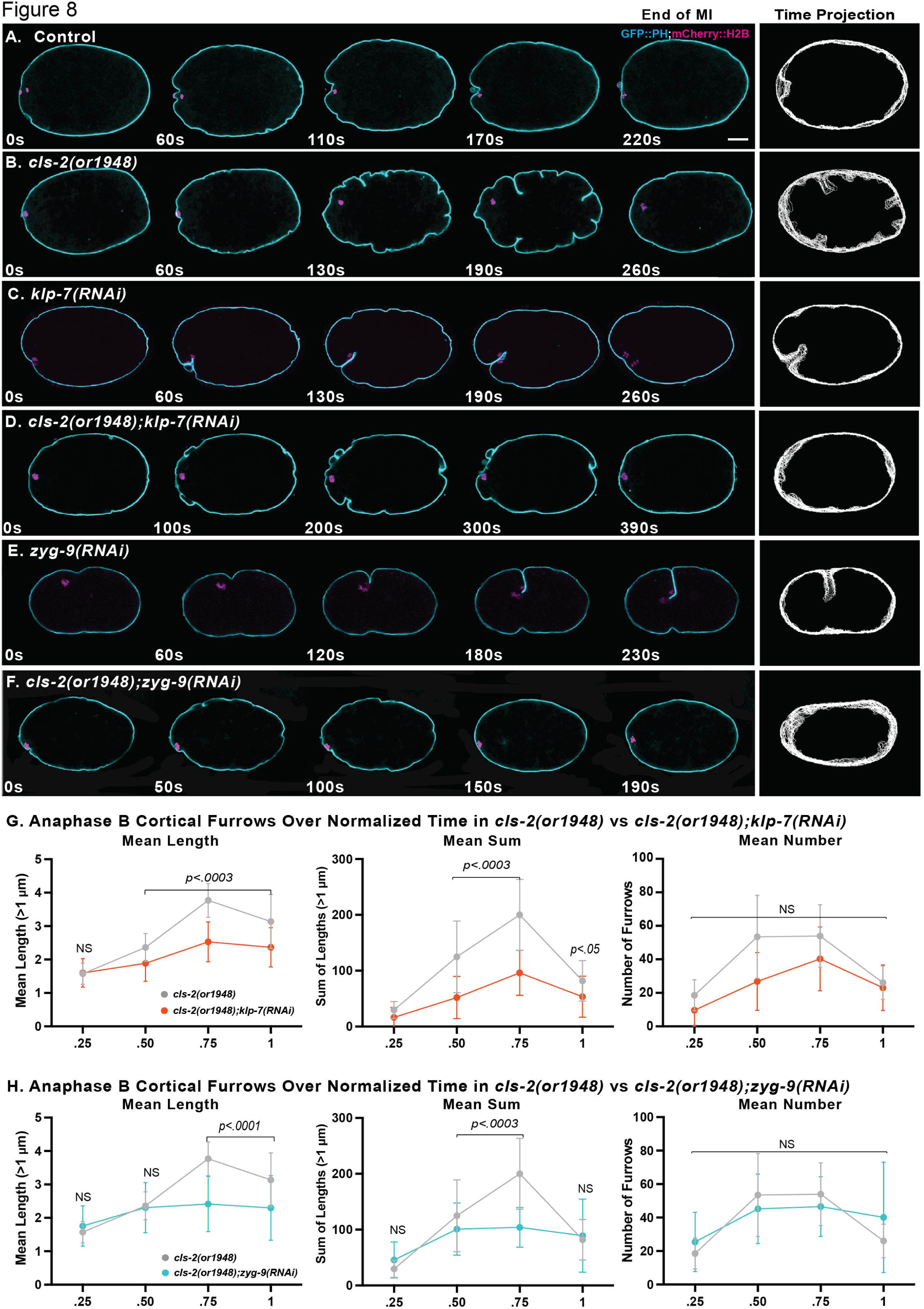
KLP-7/kinesin-13 or ZYG-9/chTOG knockdown suppresses membrane ingression in *cls-2(or1948)* oocytes. (A-F) Selected and merged focal planes during meiosis I anaphase B of live *ex utero* control and mutant oocytes expressing GFP::PH (cyan) and mCherry::H2B (magenta) to mark membranes and chromosomes, respectively. A single central focal plane is shown for the membrane, with a maximum intensity projection of 4 consecutive focal planes that encompass most of the oocyte chromosomes, which are visible at the left, anterior end of each oocyte. Time projections of single central focal planes for the membrane are shown in column at far right. (G&H) Quantification of the mean length of membrane ingressions 1μm or more in length, the mean sum of lengths, and the mean number of membrane ingressions 1μm or more in length, over normalized time in control and mutant oocytes. Note that the mean length and the mean sum of lengths, but not the mean number of ingressions, are significantly reduced in the double mutants compared to *cls-2(or1948)* single mutant oocytes. Scale bar = 10 µm.

These results further support our hypothesis that cortical microtubules limit the extent of membrane ingression during oocyte meiosis I polar body extrusion.

While the elevated microtubules levels in *cls-2 klp-7* and *zyg-9; cls-2* double mutants are correlated with the suppression of excess membrane ingression, polar body extrusion was not rescued. Instead, it was equally or more defective in the double mutants compared to the *klp-7* and *zyg-9* single mutants (Fig S3B). Furthermore, although membrane ingression in both *klp-7* and *zyg-9* mutant oocytes generally appeared reduced, we nevertheless observed deep and prominent ingressions near the oocyte chromosomes in some *cls-2, klp-7* and *zyg-9* single mutants (Figs 8C, 8E, S2, Movie 3), while prominent furrows were absent in *cls-2 klp-7* and in *zyg-9; cls-*2 double mutants (Figs 8D, 8F, S2, Movie 3), and polar body extrusion failed in nearly all single and double mutant oocytes (Fig S3B). Finally, it is notable that polar body extrusion always failed after nocodazole treatment to destabilize microtubules, but rarely failed after taxol treatment to stabilize them. Though curiously various, these results are consistent with a proper balance of cortical tension and elasticity being important for productive meiotic contractile ring dynamics in *C. elegans* oocytes.

### Microtubules influence the distribution of cortical actomyosin in C. elegans oocytes

In our previous study using *in utero* live imaging, we did not detect any differences in cortical NMY-2/non-muscle myosin dynamics during polar body extrusion in *cls-2* mutant oocytes (13). We therefore hypothesized that cortical microtubules themselves, through the biophysical properties of their tubular polymeric structure, directly confer stiffness to the oocyte cortex and thereby limit the extent of membrane ingression during polar body extrusion. Alternatively, cortical microtubules might indirectly influence membrane ingression, by signaling to cortical actomyosin in ways that we failed to detect.

To explore whether microtubules themselves, or microtubule signaling to cortical actomyosin, limits membrane ingression, we imaged the dynamics of a GFP fusion to the non-muscle myosin NMY-2 (NMY-2::GFP) in control and mutant *ex utero* oocytes.

We observed surprising and reproducible differences in the distribution of cortical NMY-2::GFP during polar body extrusion in both *klp-7* and *zyg-9* mutants, compared to control oocytes. In control oocytes, we observed the expected network of evenly distributed NMY-2::GFP foci that dissipated during anaphase B (Figs 9A, S5, Movie 6). In nearly all *klp-7* and *zyg-9* mutant oocytes, and in *cls-2 klp-7* and *zyg-9; cls-*2 double mutants, we observed more linear arrays of NMY-2::GFP foci, often but not always oriented in parallel to each other and transverse to the long axis of the oocyte (Figs 9B-E, S6, S7, Movie 6). While we have not quantified these differences, the defects appeared somewhat more severe in *zyg-9* mutant oocytes (Figs 9, S5-8). Moreover, we observed less frequent and less prominent but nevertheless similar defects in some *cls-2* mutant oocytes (Fig S8). While the spatial organization of NMY-2 foci was altered in mutant oocytes, their dissipation during anaphase B appeared similar to controls. These observations indicate that microtubules might instead, or also, act indirectly through actomyosin to limit membrane ingression (see Discussion).

**Figure 9.**
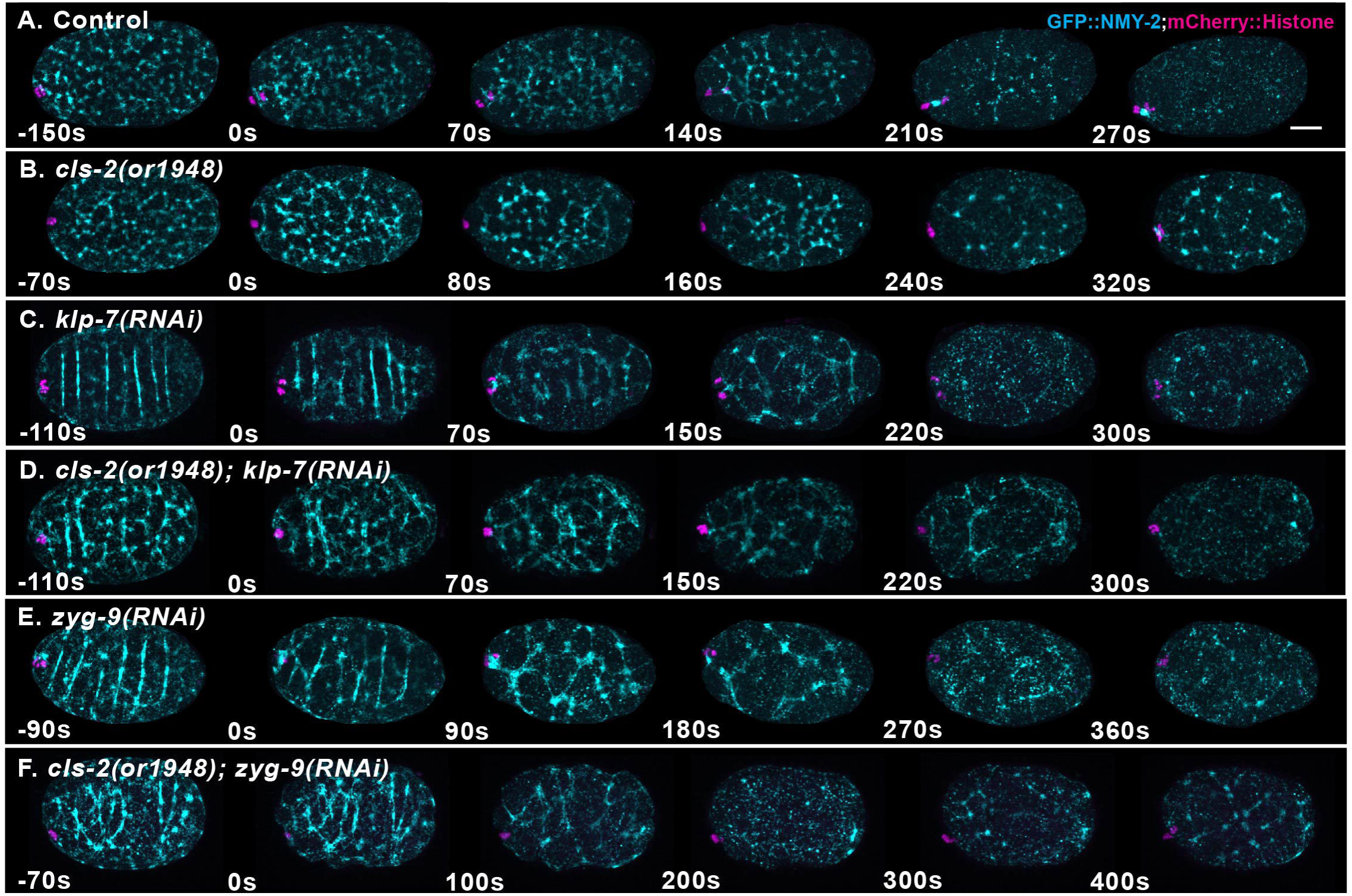
Altered levels of oocyte cortical microtubules are associated with abnormal distributions of cortical NMY-2/non-muscle myosin during *C. elegans* meiosis I. (A-F) Merged maximum intensity projections during meiosis I anaphase B of live *ex utero* control and mutant oocytes expressing NMY-2::GFP (cyan) and mCherry::H2B (magenta) to mark non-muscle myosin II and chromosomes, respectively. Maximum intensity projections of four cortical focal planes showing non-muscle myosin (cyan) are merged with maximum intensity projections of four consecutive focal planes that encompass most of the oocyte chromosomes, which are visible at the left, anterior end of each oocyte. The NMY-2::GFP foci formed extended and sometimes transversely oriented and parallel linear arrays in*klp-7* and *zyg-9* mutant oocytes. The NMY-2::GFP distribution appeared more normal in*cls-2or1948)* oocytes, but similar abnormal and extended arrays were observed in some*cls-2* mutant oocytes (see Fig S10). Scale bar = 10 µm.

After documenting defects in both cortical NMY-2 dynamics and polar body extrusion in *klp-7, zyg-9* and *cls-2* mutant oocytes, we next examined contractile ring assembly and ingression in control and mutant oocytes. In our previous study, we found that *cls-2* mutants have early defects in ring assembly or stability (13). To inspect the penetrant polar body extrusion defects in *cls-2, klp-7* and *zyg-9* mutant oocytes (Fig S3B), we used Imaris software to rotate and merge three-dimensional renderings of our NMY-2::GFP live imaging data and thereby observe contractile ring dynamics from a more oocyte end-on and ring-centric view (Figs 10, S9, Movie 7).

**Figure 10.**
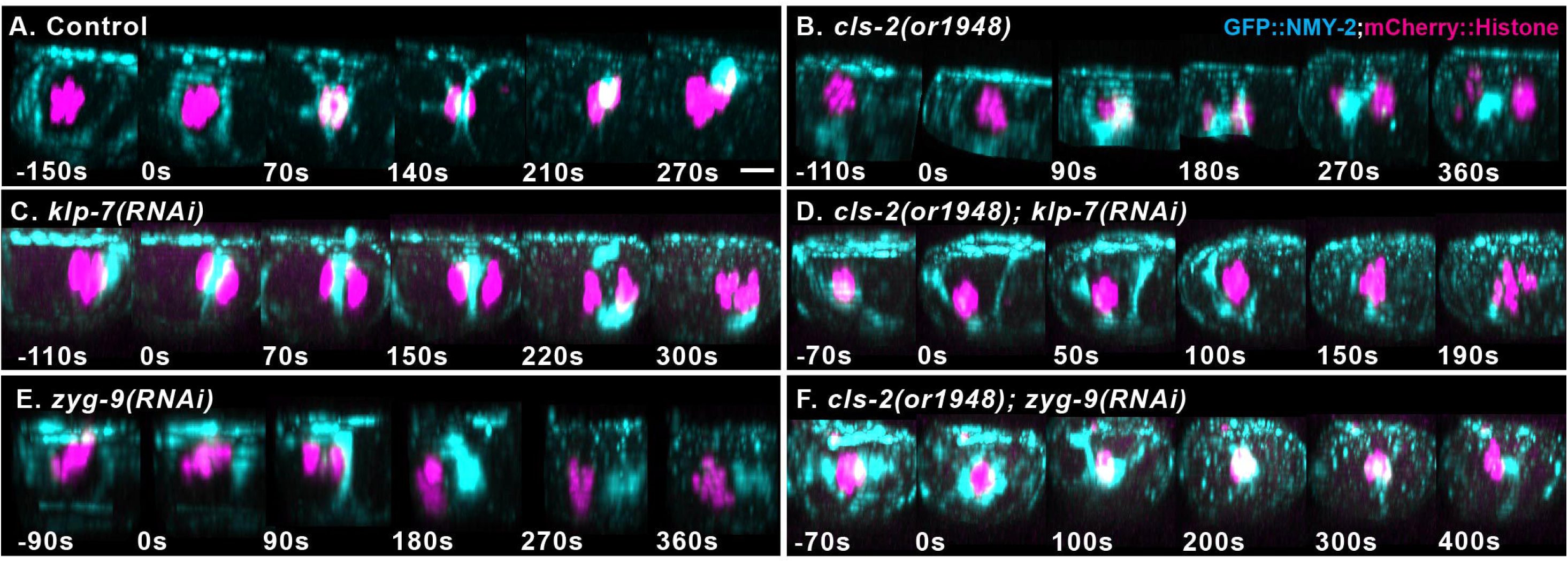
Contractile ring dynamics during meiosis I polar body extrusion are defective in*cls-2, klp-7* and *zyg-9* mutant oocytes. Merged maximum intensity projections of all focal planes after Imaris-mediated rotation to obtain ring-centric, more oocyte-end-on views of contractile ring dynamics in control and mutant oocytes expressing NMY-2::GFP (cyan) and mCherry:H2B (magenta) to mark non-muscle myosin and chromosomes, respectively. A ring forms early in anaphase B and constricts in control oocytes and but rings either are not apparent or are unstable in mutant oocytes, with NMY-2::GFP often merging into one or a few bright puncta. Scale bar = 5 um.

We observed early, similar and penetrant defects in meiosis I contractile ring assembly or stability in *cls-2, klp-7* and *zyg-9* single mutants, and in both double mutants. NMY-2::GFP foci often did not form detectable rings, and even when rings were detected they were unstable, with NMY-2::GFP moving into one or a few bright foci near the chromosomes in most oocytes and ultimately failing to extrude polar bodies (Figs 10, S3B, S9; Movie 7). We have not quantified these variable and dynamic three-dimensional contractile ring defects in detail, but depleting any one of three very different regulators of oocyte spindle and cortical microtubule stability results in early and penetrant contractile ring assembly and stability defects, highlighting the importance of microtubule regulation during *C. elegans* polar body extrusion.

## Discussion

We have shown that two subunits of an outer kinetochore sub-complex—the kinase BUB-1 and the CLASP family member CLS-2, along with the outer kinetochore scaffolding protein KNL-1, co-localize to patches distributed throughout the oocyte cortex during most of *C. elegans* meiosis I cell division. As previously documented at kinetochores, CLS-2 localization to these patches required BUB-1 and KNL-1, while BUB-1 required only KNL-1, and KNL-1 was at patches independently of both BUB-1 and CLS-2. Knocking down any one of these patch components resulted both in reduced cortical microtubule levels and excess membrane ingression throughout the oocyte during polar body extrusion. Moreover, nocodazole or taxol treatment, to destabilize or stabilize cortical microtubules, respectively, led to increased and decreased membrane ingression. Finally, genetic backgrounds with increased cortical microtubule levels suppressed the excess membrane ingression in *cls-2* mutant oocytes. These results all support our hypothesis that microtubules promote cortical stiffness and thereby limit the extent of membrane ingression throughout the oocyte during polar body extrusion. Limiting the extent of membrane ingression may be important for proper assembly and stability of the small meiotic contractile ring, which forms within and remains connected to the much larger and actively contractile cortical actomyosin network during polar body extrusion.

We hypothesize that the highly regulated distribution of cortical microtubules throughout the *C. elegans* oocyte directly stiffens the cortex to resist membrane ingression during polar body extrusion. However, because NMY-2/non-muscle myosin dynamics were altered in mutant oocytes with both elevated and decreased cortical microtubule levels, we cannot rule out the alternative and more conventional view that cortical microtubules indirectly limited membrane ingression by acting through cortical actomyosin. We nevertheless speculate that the altered cortical NMY-2 distributions in mutant oocytes with altered cortical microtubule levels are not responsible for suppressing membrane ingression, but rather are a secondary, indirect consequence of increased cortical stiffness conferred by cortical microtubules themselves (see below).

KNL-1, BUB-1, CLS-2, and other cell division proteins have been described previously as being present in roughly linear cortical structures that also are present throughout the assembling oocyte meiotic spindle. These structures previously have been referred to both as rods and as linear elements (17–20), and one component can indeed form linear filaments *in vitro* and greatly extended linear structures when overexpressed *in vivo* (33). Here we refer to these cortical structures as patches, given their less linear appearance as meiosis I proceeds, and to distinguish them from the more linear spindle-associated elements.

### CLS-2/CLASP mediates only a subset of the cortical functions promoted by KNL-1 and BUB-1

Because CLS-2/CLASP family members are known to promote microtubule stability, we anticipated that CLS-2 might mediate all functional output for the KNL-1/BUB-1/CLS-2 cortical patches, stabilizing cortical microtubules and thereby limiting the extent of membrane ingression throughout the oocyte. However, we found that while knockdown of either KNL-1 or BUB-1 increased both the number of ingressions and the mean length and the mean sum of all ingression lengths, eliminating the outermost component CLS-2 increased only the mean length and the mean sum of lengths, but not the number of ingressions. This difference is significant because we observed the more extensive requirements for KNL-1 and BUB-1 after feeding RNAi knockdowns that only partially eliminate gene function (Fig 3B; see Materials and Methods), while for CLS-2 we quantified membrane ingression in oocytes from worm homozygous for the null allele *cls-2(or1948)*. We conclude that CLS-2 mediates only a subset of the KNL-1/BUB-1/CLS-2 functions that modulate oocyte membrane ingression during polar body extrusion.

Putative KNL-1and BUB-1-dependent but CLS-2 independent factors that limit the number of membrane ingressions might, like CLS-2, stabilize cortical microtubules, but in a distinct manner that limits the number rather than extent of ingressions. Consistent with this possibility, we observed distinct distributions of elevated cortical microtubules in *klp-7* and *zyg-*9 mutant oocytes. While the resolution of our spinning disk confocal microscopy data is limited, and we have not quantified these differences, the cortical microtubules in *klp-7* mutant oocytes appear more evenly distributed, while those in *zyg-9* mutant oocytes appear more highly clustered in numerous small foci. Similarly, KNL-1 and BUB-1 might promote a microtubule stabilizing activity that limits the number of membrane ingressions, while CLS-2 promotes a different form of stability that limits only their extent. However, we also observed an increase only in the length but not the number of ingressions after nocodazole treatment to destabilize nearly all oocyte microtubules. While the altered microtubule dynamics in different mutant backgrounds might have different impacts on membrane ingression, compared to the more complete and indiscriminate reduction caused by nocodazole, the normal number of ingressions observed after nocodazole treatment argues against KNL-1 and BUB-1 acting through microtubules, independently of CLS-2, to limit ingression number.

Alternatively, KNL-1 and BUB-1 might act independently of CLS-2 to limit the number of membrane ingressions by influencing cortical actomyosin dynamics. One potential target for such regulation is the kinase CSNK-1, a negative regulator of non-muscle myosin, which when reduced in function was reported to result in large and more persistent cortical NMY-2 foci and excess membrane ingression throughout the oocyte during polar body extrusion (24). We therefore used live imaging to examine membrane and NMY-2 dynamics during polar body extrusion after RNAi knockdown of CSNK-1.

Cortical NMY-2 was upregulated and polar body extrusion usually failed, but we did not observe a statistically significant increase in either the number or length of membrane ingressions during anaphase B (Figs S3B, S10). While these results do not support a role for CSNK-1 in the regulation of ingression number, other factors that regulate cortical actomyosin might influence ingression number during polar body extrusion.

Downstream effectors that might regulate the number of membrane ingressions include targets of the BUB-1 kinase domain, and of the CENP-F-like and nearly identical coiled-coil proteins HCP-1 and -2. HCP-1/2 bridge BUB-1 to CLS-2 at kinetochores and have recently been shown to promote meiotic spindle-associated microtubule stability independently of CLS-2 (19)(16). While we do not yet know if HCP-1/2 also are present in oocyte cortical patches, further investigation of BUB-1 and HCP-1/2 may identify additional outputs that influence membrane ingression number during polar body extrusion.

### Microtubules are an integral part of the *C. elegans* oocyte cortex

Crosstalk between the microtubule and microfilament cytoskeleton is extensive (11,12), but microtubules in animal cells are nevertheless generally viewed as constituting a distinct cytoskeleton that interacts with but is not part of the actomyosin-based cortex (1,6,11). Both the contractile properties of cortical actomyosin and microfilament cross-linking influence cortical tension (4), and the cross-linking of cortical microfilaments to membrane-associated proteins by Ezrin/Radixin/Moesin (ERM) family members reduces elasticity, stiffening the cortex to promote rounding during mitosis (5–7).

While it is clear that actomyosin provides both tension and elasticity within animal cell cortices, most reported studies tested only actomyosin and not microtubule factors for requirements (5–7,36). Moreover, in one study that introduced tipless cantilever Atomic Force Microscopy as a method for assessing cortical stiffness, treatment of a non-adherent fibroblast cell line with low doses of Latrunculin A to disrupt cortical actomyosin had no significant effect on cortical stiffness (37), and in a study of T-cell migration, stabilization of microtubules by taxol, while at the same time inhibiting non-muscle myosin contractility, led to an increase in cortical stiffness (38). Though examples are limited, it seems likely that microtubules contribute to cortical stiffness other animal cell types (see below).

The results we report here, and other investigations of *C. elegans* meiotic cell division, clearly indicate that microtubules are an integral part of the oocyte cortex (18,19,28,29). Microtubules and actomyosin both are highly enriched in proximity to the plasma membrane throughout the oocyte during polar body extrusion, and the levels and organization of cortical microtubules are highly regulated. Loss of either the microtubule depolymerase KLP-7/kinesin 13, or of the TOG domain and microtubule polymerase/depolymerase ZYG-9/chTOG, significantly elevated microtubule levels throughout the oocyte cortex during meiosis I. The distinct distributions of elevated microtubules in these two mutants, and the statistically significant partial dependence on CLS-2 only in *klp-7* mutants, indicate that cortical microtubules experience extensive and complex regulation during polar body extrusion.

### Do microtubules directly or indirectly limit oocyte membrane ingression?

Our hypothesis that cortical microtubules limit membrane ingression during polar body extrusion in *C. elegans* is strongly supported by our observations that: (i) nocodazole treatment to destabilize microtubules, and mutant backgrounds that decreased cortical microtubules, increased membrane ingression throughout the oocyte cortex; (ii) taxol treatment to stabilize microtubules decreased membrane ingression; (iii) increased levels of cortical microtubules in *cls-2 klp-7* and *zyg-9; cls-2* double mutants suppressed the excess membrane ingression in *cls-2* mutant oocytes. The tubular structure of microtubules makes them relatively stiff compared to microfilaments (39), and it seems plausible that, in sufficient numbers, with appropriate length distributions and perhaps with cross-linking between themselves or with microfilaments, microtubules could themselves more or less directly stiffen the oocyte cortex. However, our results do not exclude the alternative and more conventional possibility that cortical microtubules in the oocyte indirectly suppress membrane ingression by acting through cortical actomyosin.

In support of microtubules signaling to actomyosin, we observed very similar and substantial changes in cortical NMY-2/non-muscle myosin organization in both *klp-7/*kinesin-13 and *zyg-9/XMAP215* mutants that might account for the decreased membrane ingression observed in *cls-2 klp-7* and *zyg-9; cls-*2 double mutants relative to *cls-2* single mutants. In both *klp-7* and *zyg-9* mutants, and the *cls-2 klp-7* and *zyg-9; cls-2* double mutants, cortical NMY-2 foci were less evenly distributed and instead formed prominent and often parallel linear arrays transversely oriented relative to the oocyte long axis. These abnormal NMY-2 distributions appeared similar in *klp-7* and *zyg-9* mutants, although they may be more severe in *zyg-9* mutant oocytes.

The abnormal distributions of cortical NMY-2 foci in *klp-7* and *zyg-9* mutant oocytes clearly indicate that cortical microtubules influence actomyosin dynamics. However, it is not obvious how the arrangement of cortical NMY-2 into prominent linear arrays in *klp-7* and *zyg-9* mutants, relative to the more dispersed NMY-2 foci in control oocytes, might suppress membrane ingression, rather than promote more extensive ingression in the vicinity of the linear NMY-2 arrays. Moreover, we observed similar though less prominent alterations in NMY-2 distribution in *cls-2* mutant oocytes, which have increased membrane ingression.

Alternatively, the abnormal arrays of NMY-2 foci could be a secondary consequence of altered microtubule-mediated cortical stiffness, rather than being responsible for decreased membrane ingression in *klp-7* and *zyg-9* mutants. To our knowledge, it is not known how the distribution and location of cortical NMY-2 foci correlate with sites of membrane ingression during polar body extrusion. Further investigation with live imaging to examine the spatial and temporal relationships of cortical NMY-2 foci, KNL-1/BUB-1/CLS-2 patches, sites of membrane ingression, and microtubule distribution throughout meiosis I cell division should improve our understanding of these intriguing regulatory relationships (see below).

In conclusion, the abnormal distributions of NMY-2 foci we have observed in *klp-7* and *zyg-9* mutant oocytes suggest that increased cortical microtubule levels in these mutants might indirectly suppress membrane ingression by acting through cortical actomyosin. However, these results do not rule out a direct role for microtubules in promoting cortical stiffness, and microtubules could influence cortical stiffness themselves and also act indirectly through cortical actomyosin. We favor the view that the altered distributions of NMY-2 in *cls-2, klp-7* and *zyg-9* mutant oocytes do not influence membrane ingression during polar body extrusion but rather are secondary consequences of altered, microtubule-mediated cortical stiffness.

### Additional roles for cortical microtubules in *C. elegans*

Microtubules may have additional roles as a functional component of cell cortices later in *C. elegans* embryogenesis. Astral microtubules in the one-cell zygote, and in two and four cell stage embryos, stop growing when their plus ends contact the cell cortex, with plus-end capture by complexes that include dynein mediating mitotic spindle positioning during the first few embryonic cell divisions (40). The termination of microtubule plus-end growth in early embryos requires the cortically localized GTPase EFA-6, which is no longer detectable after the 4-cell stage (41). In *efa-6* null mutants, and in wild-type embryos beyond the 4-cell stage, astral microtubules continue to elongate upon reaching the cell cortex, forming extensive networks that occupy all early embryonic cell cortices. Homozygous viable, *efa-6* null mutants produce embryos with non-lethal defects in centrosome movement and spindle rocking during the first embryonic mitosis that presumably result from the growth of microtubules throughout the one-cell cortex leading to excess dynein-mediated pulling forces (41). However, it is not known whether the ubiquitous cortical microtubules observed in wild-type embryos after the 4-cell stage are important for normal development. Cell contact can orient cell division axes in *C. elegans* embryos by modulating cortical actomyosin dynamics (42). Perhaps cortical microtubule stiffness, together with contact-mediated, actomyosin-driven orientation of division axes, are in part responsible for generating the remarkably invariant pattern of cell division axes that constitute *C. elegans* embryogenesis (43).

Another function for cortical microtubules in *C. elegans* might be to compensate for an absence of Ezrin/Radixin/Moesin (ERM) family members, which mediate the cross-linking of cortical microfilaments to membrane proteins that stiffens the cortex to promote mitotic rounding in cultured insect and mammalian cells (5–7). The *C. elegans* genome encodes only a single ERM family member, MOE-1, with limited expression in several larval cell types and limited post-embryonic requirements during larval intestinal lumen and vulval development (44). Furthermore, microtubules might contribute to cortical stiffness even in animal phyla that employ multiple ERM family members during mitotic rounding. Remarkably, in an impressively high-throughput Atomic Force Microscopy screen of all human kinases for cortical stiffness factors, the only kinase identified, other than the SILK-1 kinase that phosphorylates and activates cross-linking by ERM proteins (6,7), was BUB-1 (36). Thus BUB-1 orthologs might influence cortical stiffness even in cell types that use ERM proteins during mitotic rounding. It will be interesting to test whether CLASP orthologs also influence cortical stiffness in human and other animal cell types.

### Microtubules are required for polar body extrusion in mammals and in *C. elegans*

While actomyosin appears to play very different roles during oocyte meiotic spindle assembly and positioning in *C. elegans* compared to vertebrate oocytes (see Supplemental File S1), microtubules are important for polar body extrusion in both. Even though chromosomes, or even just DNA-coated beads can independently of microtubules induce the formation of an actin cap and a surrounding contractile ring in mouse oocytes, spindle microtubules nevertheless are required for the successful completion of polar body extrusion (45). In *C. elegans*, the requirements we have documented for CLS-2/CLASP, KLP-7/kinesin-13, and ZYG-9/chTOG, and the consequences of nocodazole and taxol treatment, all indicate that microtubules play important role(s) in polar body extrusion. Further investigation of microtubule and microfilament dynamics in both vertebrate and invertebrate oocytes is needed to better understand the similarities and differences in the mechanisms that mediate oocyte meiotic cell division across animal phyla.

### Membrane protrusion versus contractile ring ingression

1. *C. elegans* and vertebrate oocyte meiotic cell division differ substantially with respect to their extracellular environments during polar body extrusion. Mammalian oocytes produce a plasma membrane with numerous protruding microvilli throughout the surface, except over the meiotic spindle pole once it approaches the cortex (Uraji, Scheffler, and Schuh 2018). A dense actin network underlies these microvilli, which take up nutrients from surrounding follicle cells. By contrast, during meiotic cell division *C. elegans* oocytes secrete a largely impermeable eggshell that maintains the ovoid zygote shape and is closely apposed to the plasma membrane (48,49).
2. *C. elegans* and vertebrate oocyte meiotic cell division differ substantially with respect to the membrane dynamics that occur during polar body extrusion. When observed *in utero*, or *ex utero* in physiological buffer conditions, the *C. elegans* oocyte meiotic contractile ring assembles at the cell cortex, distal to and overlying the nearest spindle pole (Fig 1). Actin and myosin are depleted from the cortex within the ring (23), and the ring ingresses over one pole, driving the membrane into the oocyte before constricting midway between the two poles to ultimately pinch off a small polar body with extruded chromosomes (21,22). By contrast, in mammalian and *Xenopus* oocytes, branched actin becomes highly enriched within the contractile ring, and the membrane overlying the cortex-proximal spindle pole moves outwards, consistent with requirements for Cdc42, Arp2/3 and branched microfilament polymerization pushing out the overlying membrane (50). During this protrusion, the cortically anchored contractile ring constricts without obvious ingression into the oocyte proper.

In *C. elegans* oocytes cultured in hyperosmotic medium, a similar protrusion of the membrane overlying the cortex-proximal spindle pole has been observed during polar body extrusion (21). Furthermore, to explain the role of cortical actomyosin contractility throughout the *C. elegans* oocyte cortex during polar body extrusion, it has been hypothesized to generate hydrostatic pressure that pushes out the weakened, actomyosin-depleted cortex and associated membrane within the cortically anchored ring (23). The protruding membrane is posited to pull the tethered membrane-proximal spindle pole with it, followed by constriction of the contractile ring dividing the two chromosome sets and pinching off a polar body. In support of this model, reducing the function of the kinase CSNK-1, a negative regulator of non-muscle myosin, has been shown to result in large and more persistent cortical NMY-2 foci and excessive membrane ingression throughout the oocyte during polar body extrusion (24). In some *csnk-1* mutant oocytes, the entire spindle was extruded, consistent with excess hydrostatic pressure leading to excess membrane protrusion pulling the entire spindle into a polar body. However, as noted above, the lack of a statistically significant increase in membrane ingression number or length that we observed after CSNK-1 RNAi knockdown does not support a role for hydrostatic pressure driving membrane protrusion during polar body extrusion (Fig S10). How the *C. elegans* contractile ring dynamics observed both *in utero* and *ex utero* in balanced osmotic medium are related to the more protrusive process observed *ex utero* in hyper-osmotic medium, and the role of CSNK-1 in polar body extrusion, require further investigation.

### Coordinating multiple cortical actomyosin requirements

Instead of generating hydrostatic pressure, cortical actomyosin instead might have requirements unrelated to polar body extrusion, such that additional regulation is needed to mediate contractile ring assembly and ingression within a multifunctional cortical actomyosin network. Other possible requirements for cortical actomyosin in *C. elegans* oocytes include elongation of the roughly cuboid oocyte into an elongated oval shape during ovulation, and the effective distribution and delivery of vesicles involved in the secretion of eggshell components, with both occurring during oocyte meiotic cell division (48,49, 52). We speculate that cortical microtubules contribute to a proper balance of tension and elasticity that is required to orchestrate contractile ring dynamics within the much larger and multifunctional oocyte cortical actomyosin network. As the entire network appears to be contractile during polar body extrusion, it will be interesting to determine whether contractility is required throughout the network for other functions, or if instead contractility cannot be locally restricted to just the contractile ring during polar body extrusion. Finally, the absence of KNL-1/BUB-1/CLS-2 cortical patches during meiosis II might reflect less of a need for cortical regulation as eggshell synthesis nears completion. It will be interesting to compare cortical microtubule and actomyosin dynamics during meiosis I and II, and to determine what influence eggshell production has on polar body extrusion.

Several other *C. elegans* kinetochore components also are present in patches associated with the oocyte spindle and cortex. These include HIM-10, KNL-3, KBP-1, NDC-80, ZWL-1, SEP-1 and ROD-1 (Howe et al. 2001; Monen et al. 2005; Dumont, Oegema, and Desai 2010; Pereira et al. 2018; Bembenek et al. 2007). Presumably these all co-localize with KNL-1/BUB-1/CLS-2, but that has not been documented, and whether they influence membrane ingression or other processes during oocyte meiosis is not known. Finally, in *Drosophila,* the Ald ortholog of the conserved kinetochore-associated kinase Mps1 is present in filamentous structures during oocyte meiotic cell division, and these filaments also contain other kinetochore proteins including BUBR1 and *polo* (54). While Mps1 is not conserved in *C. elegans,* it is intriguing that similar structures with kinetochore components have been reported in *Drosophila* oocytes.

### Roles for cortical microtubules in other organisms

There are examples of other cell types in which microtubules are present at the cortex and may promote stiffness. Most notably, microtubules form a cortical band in human platelets that is required for their flattened, discoid shape (55,56). Similarly, in tissue engineering experiments designed to influence T-cell migration, chemical inhibition of both actomyosin and dynein leads to a cortical band of microtubules that modulates T-cell shape (38). In developing plant cells, aligned bundles of cortical microtubules are required for proper cell shape and morphogenesis, although these bundles may act indirectly by mediating the secretion and organization of collagen fibrils that ultimately form the rigid plant cell walls (10,57). Our discovery of a role for cortical microtubules in *C. elegans* oocytes, the general tendency in past studies of cortical stiffness to focus exclusively on actomyosin (5–7,36), the identification of BUB-1 as a stiffness factor in a cultured human cell line (36), the failure to detect a significant decrease in cortical stiffness in a human cell line after disrupting microfilaments with Latrunculin A (37), and evidence for microtubules contributing to cortical stiffness in T-cells (38), all suggest that microtubules may have more extensive roles as a functional component of animal cell cortices than has been previously appreciated.

### Limits to our analysis: optogenetics, higher resolution imaging, and spindle cues

While we hypothesize that cortical KNL-1/BUB-1/CLS-2 patches promote microtubule stability throughout the oocyte cortex, we cannot rule out the alternative possibilities that spindle-associated or other undetectable pools of these proteins are responsible for increased cortical microtubule stability. Optogenetic experiments designed to recruit KNL-1, BUB-1, CLS-2 and other possible patch components such as HCP-1/2, and a heterologous microtubule destabilizing activity called Optokatanin (58), to limited areas of the oocyte cortex may provide more conclusive evidence that localized modification of microtubule stability influences local membrane ingression independently of changes to cortical actomyosin dynamics.

Understanding how the oocyte cortical cytoskeleton functions during polar body extrusion will require more comprehensive live cell imaging studies throughout all of meiosis I and II, with higher spatiotemporal resolution. As mentioned earlier, to our knowledge it is not known how the distribution of cortical NMY-2 foci correlates with patterns of membrane ingression during polar body extrusion, and the KNL-1/BUB-1/CLS-2 cortical patches are no longer detectable by the beginning of anaphase B although they are required for normal microtubule levels throughout anaphase B. In addition, our spinning disk confocal microscopy does not provide sufficiently high spatial resolution to fully assess how cortical microtubules and actomyosin are distributed relative to each other. Recent advances in light microscopy, including the use of near Total Internal Reflection Fluorescence (TIRF) microscopy to image the cell cortex in one-cell *C. elegans* zygotes at single molecule resolution (59), greatly improve our ability to assess the dynamic molecular topography of the oocyte cortex during polar body extrusion. Higher resolution imaging will greatly improve our understanding of the spatiotemporal and functional relationships between cortical microtubules and actomyosin, and between the cortical patches of KNL-1/BUB-1/CLS-2, microtubule and NMY-2 distribution, and sites of membrane ingression during *C. elegans* polar body extrusion.

Finally, how oocyte meiotic spindle cues influence contractile ring assembly, ingression, and constriction during polar body extrusion remains poorly understood. In addition to investigating how cytoskeletal regulation throughout the oocyte cortex influences polar body extrusion, it will be important to determine how cues from the meiotic spindle and its associated chromosomes interface with cortical regulation. Understanding how the microfilament and microtubule cytoskeleton interact during the complex process of *C. elegans* polar body extrusion may prove relevant to a more general understanding of animal cell cortices, and to tissue engineering strategies.

## Acknowledgements

We thank Arshad Desai, Julien Dumont, Frank McNally, and the *Caenorhabditis* Genetics Center (funded by the NIH Office of Research Infrastructure Programs) for *C. elegans* strains, Dan Dickinson and Alexander Cartagena-Rivera for helpful discussions, and Chris Doe for sharing equipment. This work was funded by NIH grant R35 GM131749 (A.R.Q., E.S., C.-H.C., B.B.), NIH Training Grant T32HD007348-32 (A.R.Q.), and the University of Oregon Office of the Vice-President for Research (A.F.).

## Materials and Methods

### *C. elegans* strain maintenance

All *C. elegans* strains used in this study are listed in Table S1 andwere maintained at 20C on standard nematode growth medium plates seeded with *E. coli* strain OP50.

### Feeding RNAi Knockdown

All RNAi experiments were carried out by feeding *E. coli* strain HT115(DE3) induced to express double-stranded RNA corresponding to the following genes: *klp-7, zyg-9, cls-2, bub-1, knl-1,* and *csnk-1.* First bacteria clones were picked from an RNAi library and grown on LB Agar plate with Ampicillin (Kamath et al., 2003). Each bacterial clone was miniprepped using the Qiagen kit for subsequent sequence confirmation. Hypochlorite synchronized L1 larvae were grown on standard nematode growth medium plates, washed with M9 three times and then plated on the induced RNAi plates and grown at 20C until imaging. The feeding times were chosen such that if treatment were extended for an additional 6 more hours, 90% or more of the adult worms became sterile. For *klp-7, zyg-9, bub-1, and cls-2* worms were fed for 48-52 hours. For *csnk-1,* worms were fed for 72 hours (Flynn et al., 2017, Panbianco et al., 2008). For *knl-1* RNAi, L1 larvae were grown until L4 stage and placed onto *knl-1* RNAi plates for 35-40 hours.

### Nocodazole and taxol treatment

Young adult worms were dissected to release oocytes in 1ul egg salt buffer (118mM NaCl, 48mM KCl, 2mM CaCl2, 2mM MgCl2, and 0.025 mM of HEPES, filter sterilized before HEPES addition) containing 2% DMSO, 20 ug/ml nocodazole or 200nM taxol, or DMSO only for control. The coverslip with oocytes were placed over 3% agarose pad with egg salt buffer containing 1% DMSO, 10 ug/ml nocodazole, or 100 nM taxol then used immediately for imaging.

### Image acquisition

All imaging was carried out using a Leica DMi8 microscope outfitted with a spinning disk confocal unit–CSU-W1 (Yokogawa) with Borealis (Andor), dual iXon Ultra 897 (Andor) cameras, and a 100x HCX PL APO 1.4–0.70NA oil objective lens (Leica). (Molecular Devices) imaging software was used for controlling image acquisition. The 488nm and 561nm channels were imaged simultaneously with 1um Z-spacing.

*Ex utero* live imaging used the following parameters: 1μm Z spacing, 16 focal planes, 100ms exposure, 10s interval. Imaging was carried out by dissecting worms in 3ul egg salt buffer (118mM NaCl, 48mM KCl, 2mM CaCl2, 2mM MgCl2, and 0.025 mM of HEPES, filter sterilized before HEPES addition) on a coverslip before mounting onto a 2% agarose pad (diluted in egg salt buffer) on a microscope slide.

*In utero* live imaging used the following parameters: 1 μm Z spacing, 20 μm stacks, 80s exposure, 20s interval. Imaging was carried out by placing adult worms in 1.5 μl of M9 mixed with 1.5 μl of 1 μm polysterene microspheres (Polysciences Inc.) on a coverslip which was mounted carefully onto a 5% agarose pad.

### Image processing

General image processing was performed with ImageJ/Fiji software (National Institutes of Health). First, raw images were merged for the red/green channels then cropped to a 512×512 frame in order to make montages for each figure starting from Anaphase B to the end of anaphase B/end of meiosis I. Timing of each oocyte relied on the three-dimensional projection made by using Imaris software (Bitplane). Beginning of anaphase B was determined by the chromosome channel where the chromosomes were most compact during anaphase, as described elsewhere (REF). The end of anaphase B/meiosis I was the first timepoint when oocyte chromosomes began to decondense after attempts at polar body extrusion (Schlientz, 2020). For quantification analysis and figures, areas around oocytes were cropped out. Quantification of cortical microtubules and ingressions were analyzed in a normalized anaphase B time window to account for differing lengths of anaphase B duration in each genotype (Fig S3A). Occytes were normalized by dividing the time intervals into four equal parts.

### *C. elegans* Oocyte Cortical Microtubule Patch Analysis Pipeline

Cortical microtubule patch quantification was carried out by initial processing using ImageJ/Fiji software followed by Imaris Software 9.8.2 (Bitplane). Macros available upon request.

*Fiji/ImageJ was used to isolate locally strong TBB-2 signal and histone H2B signal.* The Fiji/ImageJ processing step produces a total of 5 channels: Channel 1 – Original Histone H2B (chromosome); Channel 2 – Original TBB-2 (microtubule); Channel 3 – H2B with Median, Gaussian, and Maximum Filters; Channel 4 – Tubulin with Remove Outliers, Subtraction and Median Filters; Channel 5 – Mask Segmentation (from the maximum intensity Z projection of Channel 2).

Once the raw image files for the GFP and mCherry channels were merged, the histone H2B and TBB-2 channels were filtered to isolate locally strong signals from each channel, and eliminate background. First, the original histone H2B channel was duplicated and processed with the following filters: Median filter with radius 5 pixels, Gaussian Blur, with sigma radius of 10 pixels, and Maximum filter with radius 5 pixels; refer to macro titled ‘MedianGaussianBlurMaximum.’ These filters expand the isolated H2B signals to encompass the adjacent tubulin signals. This expansion was used to make surfaces in Imaris to eliminate sperm associated signals. This result were later be merged with the original merged image file as Channel 3.

Second, the TBB-2 channel was processed with the remove outliers Fiji function (Plugins/Integral Image Filters/Remove Outliers) for background subtraction to isolate the cortical microtubule patch signals visible in the GFP::TBB-2;mCherry::H2B associated strains with the following parameters: Block radius X 20 pixels, Block radius Y 20 pixels, and standard deviation .90 pixels. The result produced was subsequently used to model the background to subtract from the original TBB-2 channel using Fiji’s Process/Image Calculator function. Furthermore, to remove small, noisy, or sparse signals the median filter was applied using a radius of 5 pixels. The final result produces an image of isolated, locally strong signal intensity, which includes cortical microtubule patches, sperm associated microtubules, spindle microtubules, and signals outside the oocyte. Refer to macro titled ‘DuplicateAndRemoveOutlier.ijm’ This result was later merged with the other created channels and original merged file as Channel 4.

Next, the original raw microtubule channel was duplicated again and a maximum intensity z-projection function was run to prepare for hand segmentation. For each timepoint, the region outside of the oocyte was hand segmented and cropped using Fiji’s Edit/Clear Outside function. Afterward, the movie was converted to an 8-bit format. Refer to a macro titled ‘TurnTo1and016bit.ijm.’ Run the multiply function with a value of 255 followed by the divide function with a value of 255. Then convert back to 16-bit.

This process converted the hand segmented picture into an 16-bit binary picture with values of 1 inside the ROI and values of 0 outside the ROI. This created a mask of the segmented oocyte so that Imaris processing (described below) could automatically make a surface roughly surrounding the oocyte without having to consider objects outside of the oocyte.

Next we ran Fiji’s Image/Stack/Tools/Concatenate function of this mask 4 times. This was to make a hyper stack with 16 slices in the z dimension from the mask derived from the Max projected image with only 1 z slice. Following this, run the function ‘stack to hyperstack’ with the following parameters: set to xyczt, channels – 1, slice – 16, and frames – number of frames in the original image file. The end product will be a mask of the segmented oocyte with 16 stacks in the z dimension. This result will later be merged as Channel 5.

Finally, refer to a macro titled ‘MarkZandNextTimepoint.ijm’. This step was to mark the Z plane of the top surface of the oocyte. Duplicate the original TBB-2 channel. Starting from time point 1, draw a rectangle ROI around the oocyte of the outermost (cortical) z plane position of the oocyte. Run the macro for each timepoint, adjusting the z plane to the outermost cortical surface position that first shows the tubulin signal. This created a mask that has extremely high intensity values than the surrounding region and was automatically recognized by the ‘Correct 3D Drift’ plugin described below.

This result was later merged with the other created channels as Channel 6.

Run the merge function for all the newly created channels and the original Histone H2B and Microtubule TBB-2 channels:

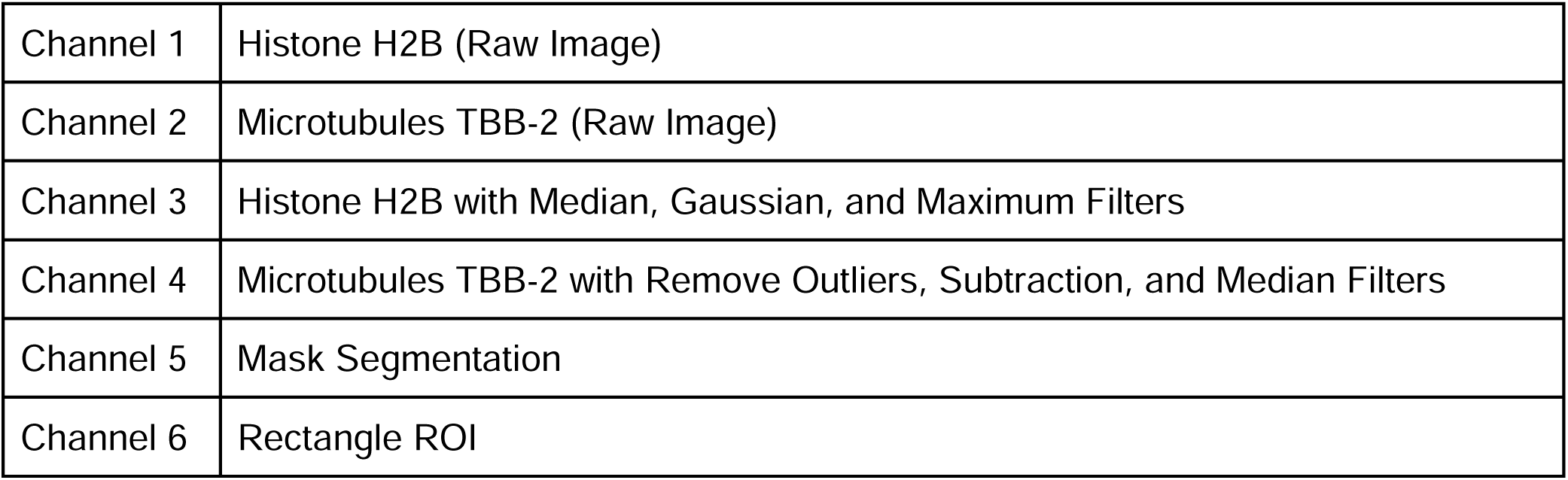

|  |  |
| --- | --- |
| Channel 1 | Histone H2B (Raw Image) |
| Channel 2 | Microtubules TBB-2 (Raw Image) |
| Channel 3 | Histone H2B with Median, Gaussian, and Maximum Filters |
| Channel 4 | Microtubules TBB-2 with Remove Outliers, Subtraction, and Median Filters |
| Channel 5 | Mask Segmentation |
| Channel 6 | Rectangle ROI |

In order to correct for 3D drift, channel 6 was used with the following parameters:

|  |  |
| --- | --- |
| Channel Registration | Channel 6 |
| Only consider pixels with values larger than 0 | Set to 0 |
| Lowest z plane to take into account | 1 |
| Highest z plane to take into account | 16 |
| Max shift x(pixels) | 10 |
| Max shift y(pixels) | 10 |
| Max shiftzy(pixels) | 90 |

This will correct drift based off of the channel 6 ROI. Among the resulting z-slices, slices with the oocyte signal are duplicated leaving z-slices outside the oocyte. At this time only channels 1-5 are duplicated because channel 6 will not be used hereafter. This image file with 5 channels will be used for Imaris (Bitplane) for further segmentation, drift correction and to make surfaces corresponding to the cortical microtubule signals Imaris 9.8.2 (Bitplane)

Surfaces 1 - Oocyte

| Parameter | yes/no | value |
| --- | --- | --- |
| Segment only a region of interest | no | N/A |
| Track Surfaces (Over Time) | yes | N/A |
| Classify Surfaces | no | N/A |
| Object-Object Statistics | yes | N/A |
| Source Channel | N/A | 5 |
| Smooth (Surface Detail) [um] | yes | 0.294 |
| Thresholding (Absolute Intensity) | yes | Automatic |
| Filters | yes | 6.34 e4 voxels |
| Edit Surfaces | no | N/A |
| Tracking Algorithm | N/A | Autoregressive motion? |
| Tracking Max Distance | N/A | 50um |
| Tracking Max Gap Size | yes | 0 |
| Fill gaps with all detected objects | no | N/A |
| Filter Tracks | no | N/A |

Open the merged dataset containing the generated channels 1 through 5 in Imaris. Create a new surface and refer to the custom creation parameter ‘Oocyte.’ Under algorithm settings mark track surfaces and object-object statistics, indicate source channel 5 (name not specified), smooth setting, thresholding absolute intensity, and surface detail defined at 0.294 um. Result should create a gray surface that encompasses the oocyte. Under the edit icon in the properties arena, mask all - channel 2 setting was used with the set voxel outside surface to 0. This created a masked channel 2.

Surfaces 2 - Oocyte Refine

Deselect Surfaces 1, and select all channels except channel 1 and masked channel 2 Create a new surface and refer to the custom creation parameter ‘Oocyte Refine.’ This step was for further drift correction. The following settings were applied: Track surfaces, source channel 6 (also referred to as masked channel 2), threshold, algorithm autoregressive motion with parameters set to max distance 50 um and gap size 0. To correct drift, select all tracks and connected objects, correct image and all objects, translational and rotational drift, and include the entire result. Select channel 4 (the channel created in ImageJ) and masked channel 2. Under the edit icon indicate ‘Mask All’ for the channel 4 selection, with mask settings as constant inside/outside set voxels outside surface to 0, and applied to all time points. This results in a new masked channel 4.

Surfaces 3 - Nucleus Expand

Create new surface 3 and refer to the custom creation parameter ‘NucleusExpand.’ Set the source channel 3 with smoothing surface details .294 um and Absolute Intensity thresholding. Under the edit tab within properties arena set ‘Mask All’ with the following parameters: Channel Selection Channel 7 - Masked Channel 4, Constant inside/outside set voxels inside surface to 0 um, and apply to all time points. This creates a “Masked Masked channel 4”. This was used to eliminate the sperm associated TBB-2 signal from the isolated TBB-2 patch signals.

Surfaces 4 - Quantification of cortical microtubule patches

Select Channel 1, 2, 4, and Masked Masked Channel 4 under the display adjustment. Create new surfaces 4 and refer to creation parameters titled ‘CMPExUteroR2.’ Select track surfaces, classify surfaces, and object-object statistics. Next select the source channel 8 - Masked Masked Channel 4 with smoothing surfaces detail set to 0.294 um, and thresholding background subtraction Diameter of largest sphere which fits into the object 10 um. The result will show the filtered cortical microtubule surfaces. Manual deletion of surfaces outside of the oocyte were done for each time point. Set algorithm to autoregressive motion with max distance parameter 5 um and max gap size 3. From this point each of the surfaces will be defined and categorized by their mean intensity.

Deselect surfaces 4 and select surfaces 3. Under the edit tab select mask all and channel 6 - masked channel 2. Under mask settings select constant inside/outside and set voxels outside surface to 0 um. This step was used to make a surface that encompasses the spindle. This creates a Masked Masked channel 2.

Surfaces 5 - Spindle

Deselect all display adjustment channels except for Channel 1 and Masked Masked channel 2. Create new surfaces 5 and choose spindle parameters with source channel 9 - masked masked channel 2. Use the recommended threshold (absolute intensity) value. This creates a single surface of where the spindle was located through all time points. Furthermore, we ran the autoregressive motion algorithm with parameters set to max distance 50 um and max gap size 0.

From here the statistics for surfaces 4 that define cortical microtubule patches can be exported. CMPs of which the distance from the sindle surface is larger than 0.1 um are subjected to statistical analysis. The voxel intensity cut off for detection of CMP surface was 150 of channel 4. For classification, CMPs having mean intensity of channel 2 lower than 2303 were classified as weak, between 2303 and 2999 were classified as medium, higher than 2999 were classified as strong. These thresholds are set based on control movies, and were used for all other genotypes. Below shows histograms of three control movies with the cutoffs:

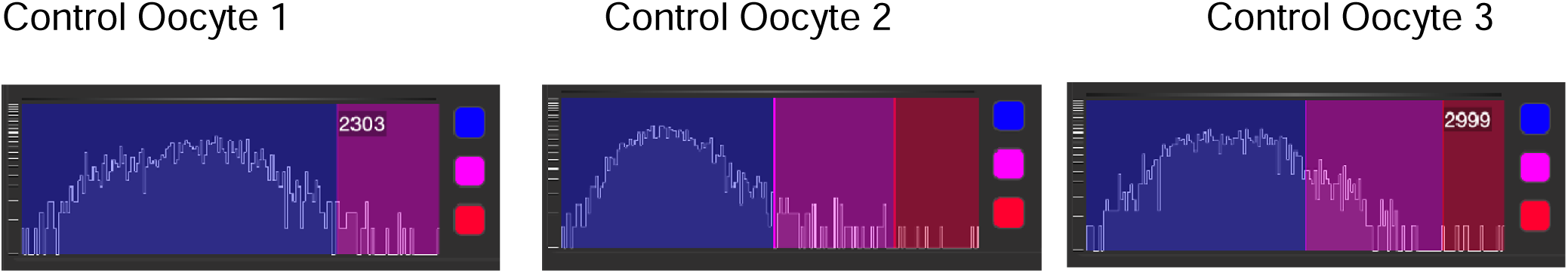

The histograms are based on the mean intensity of all surfaces. Each surface represents a single cortical microtubule patch (CMP. The histograms are automatically made in Imaris throughout the course of image processing. The classification threshold between class for weak and medium 2303 was set to separate the left most mountain, best represented in Control Oocyte 1 and Control Oocyte 2. Control oocyte 1 was a darker example of the three presented, and shows a high frequency of weak CMPs. Control oocyte 2 was a brighter example, and the thresholding cutoffs are represented with the thresholding covering the whole mountain within the weak CMP range, and lower frequency for medium, and strong CMPs. The weak class does not cover the whole left-most mountain in control oocyte 3, a very bright example, and CMPs with high intensity in the mountain are classified as medium. But the bright patches outside the mountain are classified as the strong class with the threshold 2999.

### *C. elegans* Global Cortical Ingression Analysis Pipeline

Manual segmentation was performed in FIJI to isolate dissected oocytes from adjacent dissected biological objects and subsequent automated segmentation and measurements were performed using python3 (skimage, scipy, tifffile, numpy, pandas, and matplotlib libraries). The algorithm developed was motivated by the following (make a legend or a descripton).

1. First, the oocyte cortex was modeled as a contour (Figure 1).
2. To measure the length of the ingressions, the convex hull of the cortex contour was used to establish a measurement reference to the ingressions. Given a set of points the convex hull was the subset of points that form lines which, when connected, enclose all other points (Figure 2).
3. In the absence of ingressions, the contour and convex hull completely overlap (Figure 3). When ingression begins, the convex hull reference lines emerge from the ingressing contour (Figure 4).
4. Measurements were made piece-wise between adjacent intersection points of the convex hull and contour. The ingress measurement could then be transformed from 2D coordinate space to 1D distance space by finding the distance from the contour at any point to the reference line of the convex hull. Python s scipy library was used to find peaks in the distance spac and any peak distance longer than 0.441 m (3 pixels) from the convex hull was recorded as a ingress length.
5. Distances were calculated using the following equation. and are the adjacent intersection points between the convex hull and contour, was any point along the between and (Figure 5). These distances are calculated by using the length of the cross product between ________vectors and, normalized by the length of vector :

**Figure 1:**
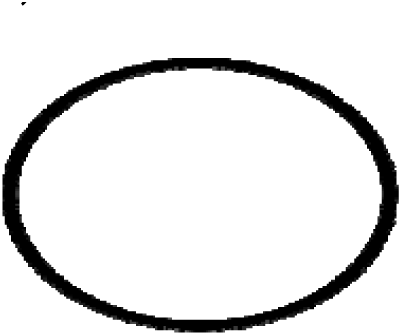
Contour model of an oocyte cortex without ingressions.

**Figure 2:**
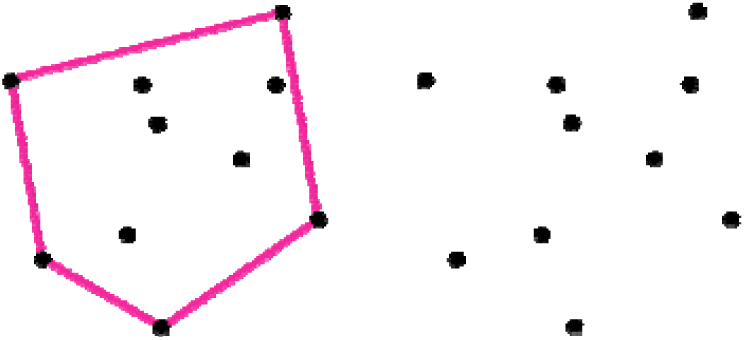
Visual description of a convex hull (magenta).

**Figure 3:**
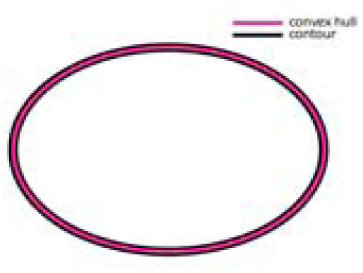
The contour and convex hull model of the oocyte cortex without ingressions.

**Figure 4:**
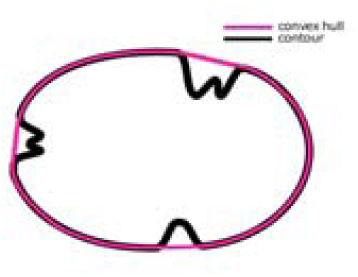
Difference between the contour model and convex hull when ingression occurs.

**Figure 5:**
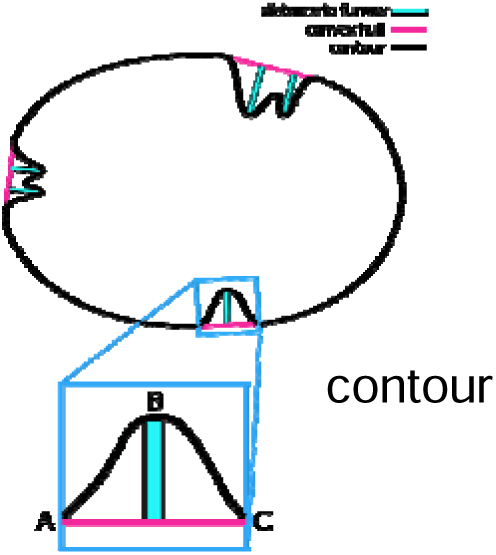
Schematic for the calculation of ingress lengths from measuring the distances from the line joined by intersection points and to any point along the contour.

### Algorithm

1. Movies were first hand-segmented using Fiji s elliptical brush tool to remove adjacent oocytes and non-relevant biological material.
2. A median filter using a pixel kernel was performed on each frame followed by intensity-based threshold using Yen s algorithm to produce a binary image of the oocyte cortex (Yen JC, Chang FJ, Chang S. A new criterion for automatic multilevel thresholding. IEEE Trans Image Process. 1995;4(3):370-8. doi: 10.1109/83.366472. PMID: 18289986.).
3. When the contour ingresses and appears to form a line (narrow, deep ingress), the contour model fails to fully capture the length of the ingression. To account for this, the following additional steps were added to the segmentation process to minimally accentuate the morphology in a uniform way (Figure 6).

a. Once the image iwas binarized after Yen thresholding, it was then dilated by 1 pixel and inverted.
b. The inverted image has now two non-zero objects, the interior of the cortex and the exterior which was shaped by the image edges.
c. The exterior, non-zero object touching the image edge was removed leaving the interior as the basis for the contour modeling. This interior retains the extent of these narrow, deep ingress.
Once the images are fully segmented, the convex hull and contour are calculated and their intersection points found. These points serve as the basis of the analysis above.
Ingressions within close proximity to chromosome location (also referred to as spindle-associated ingressions) were excluded in the analysis. To discount these ingressions, the maximum intensity projection of the chromosome channel was used as a positional mask against the cortex channel.

d. Yen thresholding was used on the maximum intensity projection of the chromosome channel along the focal axis (z).
e. The pixel locations of the binary chromosome objects were stored for every time point.
f. Ingress peak positions found to be within 3.23_µ_m (22 pixels) of the chromosome objects were classified as being spindle-associated ingressions and not included into the final analysis. The constraint of 22 pixels was measured to be the largest possible search radius from chromosome edges to potential ingress edges to exclude main ingressions that included multiple ingression points in their entirety. Once algorithm was completed, ingression lengths were exported into an excel sheet. Ingression lengths are filtered out for any ingressions greater than 1 micron and any ingressions in close proximity to chromosomes. Mean length, mean sum, and mean number are calculated in the normalized anaphase B time window.

**Figure 6:**
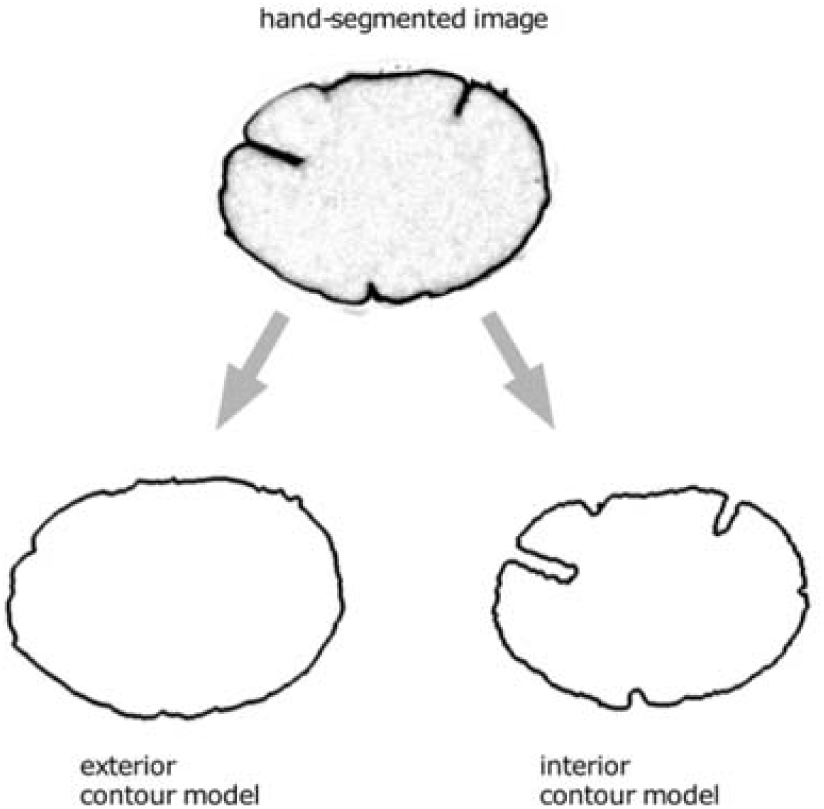
Inverting the intensity-based segmentation of the cortex allows for a better contour model of the ingressions. The contour on the bottom-left uses the intensity-based threshold of the cortex fluorescence as the input for the contour model (exterior contour model), but poorly captures the narrow ingressions. Instead we use the cortex intensity as a border and the inverted interior as the input for the contour model (bottom-right).

### Statistics

P-values comparing distributions for microtubule and ingression quantification were calculated using the Mann-Whitney U-test. Statistical analysis was performed using Microsoft Excel (Microsoft) and Prism 9 (GraphPad Software) and graphs were made in Prism 9 (see Supplemental Files S2, S3).

## Supplemental Information

**Figure S1. CLS-1, BUB-1 and KNL-1 cortical patches are present only during meiosis I, not during meiosis II.** Maximum intensity projections of all focal planes during meiosis I in live ex utero oocytes expressing mCherry::H2B (magenta) to mark chromosomes and GFP (cyan) fusions to CLS-2 (A), BUB-1 (B), and KNL-1 (C). All three proteins were associated with the oocyte spindle and chromosomes, and were present in cortical patches, during meiosis I, but were associated only with the spindle and chromosomes, and not present in cortical patches, during meiosis II.

**Figure S2. Anaphase B membrane ingression in control and mutant oocytes.**

Time projections of single central focal planes throughout meiosis I anaphase B from six ex utero oocytes of each genotype or condition, all expressing GFP::PH (cyan) and mCherry::H2B (not shown) to mark oocyte membranes and chromosomes, respectively.

**Figure S3. Cell division defects and computational quantification of membrane ingression during *C. elegans* oocyte meiosis I.**

(A) Anaphase B duration in oocytes of each genotype or condition, with mean and standard deviation indicated. (B) Bar graphs indicating number of embryos scored and percent of each genotype or condition in which polar body extrusion after meiosis I was successful, with some mCherry::H2B signal detected in an intact polar outside of the oocyte after the completion of meiosis I. (C) Schematic of a convex hull and contour model to measure furrow lengths in *C*. elegans oocytes. Defects in the contour are shown in middle oocyte and are quantified using a distance formula (see Materials and Methods). (D) Single focal planes of live control and mutant oocytes strains that express GFP::PH (white) and mCherry::H2B (not shown) processed for scoring ingression. Pink dots indicate furrows that are computationally counted and measured in length (see Materials and Methods).

**Figure S4. Imaris quantification of cortical microtubule patches (CMPs) in control and *cls-2(or1948) oocytes*.**

(A,B) Merged maximum intensity projections of *ex utero control* and *cls-2(or1948)* oocytes that express GFP::TBB-2 (white) to mark microtubules, and mCherry::H2B to mark chromosomes (not shown). Top rows for control and *cls-2(or1948)* oocytes show GFP::TBB-2 (white); middle rows show color-coded Imaris surfaces, based on our cutoffs for weak (blue), medium (purple), and strong (red) CMPs (see Materials and Methods); bottom rows show merges of GFP::TBB-2 and Imaris surfaces. (C) Quantification of weak, medium and strong CMPs in single control and *cls-2(or1948)* oocytes during anaphase B. The *cls-2(or1948)* oocyte shown has mostly weak CMPs, with a loss of medium and strong CMPs throughout anaphase B.

**Figure S5. Cortical non-muscle myosin distribution in control oocytes.**

Merged maximum intensity projections during meiosis I anaphase B of five *ex utero* control oocytes expressing NMY-2::GFP (cyan) and mCherry::H2B (magenta) to mark NMY-2/non-muscle myosin II and chromosomes, respectively. Maximum intensity projections of four cortical focal planes showing non-muscle myosin are merged with maximum intensity projections of four consecutive focal planes that encompass most of the oocyte chromosomes, which are visible at the left, anterior end of each oocyte. A roughly even but variable and dynamic network of cortical NMY-2::GFP foci persists throughout most of anaphase B until dissipating near the end.

**Figure S6. Cortical non-muscle myosin distribution in cls-2 mutant oocytes.**

Merged maximum intensity projections during meiosis I anaphase B of five *ex utero cls-2(or1948)* oocytes expressing NMY-2::GFP (cyan) and mCherry::H2B (magenta) to mark NMY-2/non-muscle myosin II and chromosomes, respectively. Maximum intensity projections of four cortical focal planes showing non-muscle myosin are merged with maximum intensity projections of four consecutive focal planes that encompass most of the oocyte chromosomes, which are visible at the left, anterior end of each oocyte. Note that abnormal linear arrays of NMY-2::GFP foci are present in some cls-2 mutant oocytes (arrowheads).

**Figure S7. Cortical non-muscle myosin distribution after KLP-7 knockdown.**

Merged maximum intensity projections during meiosis I anaphase B of five *ex utero klp-7(RNAi)* oocytes expressing NMY-2::GFP (cyan) and mCherry::H2B (magenta) to mark NMY-2/non-muscle myosin II and chromosomes, respectively. Maximum intensity projections of four cortical focal planes showing non-muscle myosin are merged with maximum intensity projections of four consecutive focal planes that encompass most of the oocyte chromosomes, which are visible at the left, anterior end of each oocyte.

**Figure S8. Cortical non-muscle myosin distribution after ZYG-9 knockdown.**

Merged maximum intensity projections during meiosis I anaphase B of five *ex utero zyg-9(RNAi)* oocytes expressing NMY-2::GFP (cyan) and mCherry::H2B (magenta) to mark NMY-2/non-muscle myosin II and chromosomes, respectively. Maximum intensity projections of four cortical focal planes showing non-muscle myosin are merged with maximum intensity projections of four consecutive focal planes that encompass most of the oocyte chromosomes, which are visible at the left, anterior end of each oocyte.

**Figure S9. CSNK-1 depletion results in abnormal cortical NMY-2::GFP dynamics and polar body extrusion defects, but not in excess membrane ingression, during meiosis I anaphase B.**

(A,B) Selected and merged focal planes during anaphase B from *ex utero* control and *cls-2(or1948)* oocytes expressing GFP::PH (cyan) and mCherry::H2B to mark membrane and chromosomes, respectively. A single central focal plane is shown for the membrane, with a maximum intensity projection of 4 consecutive focal planes that encompass most of the oocyte chromosomes, which are visible at the left, anterior end of each oocyte. Time projections of all anaphase B single central focal planes during meiosis I are at far right. (C) Time projections of single central focal planes from *csnk-1*(*RNAi*) oocytes expressing GFP::PH (white), organized by polar body extrusion phenotype: (i) large polar body, (ii) some chromosome-associated membrane ingression but failed extrusion, and no ingression detected. (D) Quantification of polar body extrusion phenotypes. In the *csnk-1*(*RNAi*) oocytes in which polar body extrusion was scored as successful (see Materials and Methods), the polar bodies were abnormally large. In the taxol treated oocytes in which polar body extrusion was scored as successful, the polar bodies appear normal in size (for an example, see Fig 6). (E) Quantification of membrane ingressions over normalized anaphase B time in control and *csnk-1*(*RNAi*) oocytes; no significant difference in mean length, mean sum of lengths, or mean ingression number were observed in *csnk-1* mutants compared to control oocytes. (F&G) Merged maximum intensity projections during meiosis I anaphase B of control and *cls-2(or1948)* oocytes expressing NMY-2::GFP (cyan) and mCherry::H2B (magenta) to mark NMY-2/non-muscle myosin II and chromosomes, respectively. Maximum intensity projections of four cortical focal planes showing non-muscle myosin are merged with maximum intensity projections of four consecutive focal planes that encompass most of the oocyte chromosomes, which are visible at the left, anterior end of each oocyte. The NMY-2::GFP foci in *csnk-1*(*RNAi*) oocytes appeared larger and less numerous, and they persisted at the cortex throughout anaphase B instead of dissipating as in control oocytes.

**Figure S10. Contractile ring dynamics during meiosis I polar body extrusion are defective in control and mutant oocytes.**

Projections of all focal planes after Imaris-mediated rotation to obtain ring-centric, more oocyte-end-on views of contractile ring dynamics in four each of control and mutant oocytes expressing NMY-2::GFP (cyan) and mCherry:H2B (magenta) to mark NMY-2/non-muscle myosin II and chromosomes, respectively, in control and mutant oocytes (see Materials and Methods).

**Movie 1. KNL-1/BUB-1/CLS-2 cortical patch dynamics during meiosis I and II.**

Full maximum intensity projections of all focal planes from (top row) in utero oocytes from nuclear envelope breakdown to end of meiosis I, and (bottom row) ex utero oocytes from meiosis I metaphase to end of meiosis II.

**Movie 2. Co-localization of KNL-1, BUB-1 and CLS-2 during meiosis I.**

Merged maximum intensity projections of four cortical focal planes and four consecutive focal planes that encompass most of the oocyte chromosomes.

**Movie 3. Membrane dynamics in control and mutant oocytes during meiosis I anaphase B**

Merged single central focal plan for membrane and four consecutive focal planes that encompass most of the oocyte chromosomes.

**Movie 4. Cortical microtubules in control and mutant oocytes during meiosis I anaphase B**

Merged maximum intensity projections of four cortical focal planes and four consecutive focal planes that encompass most of the oocyte chromosomes.

**Movie 5. Imaris spots quantification of microtubules in control and cls-2 mutant oocytes**

Projections of all focal planes after Imaris-mediated rotation.

**Movie 6. Cortical NMY-2/non-muscle myosin distribution in control and mutant oocytes during meiosis I anaphase B**

Merged maximum intensity projections of four cortical focal planes and four consecutive focal planes that encompass most of the oocyte chromosomes.

**Movie 7. Imaris-rotated views of contractile ring dynamics in control and mutant embryos during meiosis I anaphase B.**

Projections of all focal planes after Imaris-mediated rotation

**Table S1.** *C. elegans* strains used in this study.

**Supplemental File S1.** Phylogenetic diversity in oocyte meiotic cell division mechanisms.

**Supplemental File S2.** Raw data for calculation of cortical microtubule patch p values.

**Supplemental File S3.** Raw data for calculation of membrane ingression p values.

## References

1. Chugh P, Paluch EK. The actin cortex at a glance. Journal of Cell Science. 2018 Jul 15;131(14):jcs186254.

2. Koenderink GH, Paluch EK. Architecture shapes contractility in actomyosin networks. Current Opinion in Cell Biology. 2018 Feb;50:79–85.

3. Svitkina TM. Actin Cell Cortex: Structure and Molecular Organization. Trends in Cell Biology. 2020 Jul;30(7):556–65.

4. Chugh P, Clark AG, Smith MB, Cassani DAD, Dierkes K, Ragab A, et al. Actin cortex architecture regulates cell surface tension. Nat Cell Biol. 2017 Jun 1;19(6):689–97.

5. Maddox AS, Burridge K. RhoA is required for cortical retraction and rigidity during mitotic cell rounding. Journal of Cell Biology. 2003 Jan 20;160(2):255–65.

6. Carreno S, Kouranti I, Glusman ES, Fuller MT, Echard A, Payre F. Moesin and its activating kinase Slik are required for cortical stability and microtubule organization in mitotic cells. Journal of Cell Biology. 2008 Feb 25;180(4):739–46.

7. Kunda P, Pelling AE, Liu T, Baum B. Moesin Controls Cortical Rigidity, Cell Rounding, and Spindle Morphogenesis during Mitosis. Current Biology. 2008 Jan;18(2):91–101.

8. Ambrose C, Allard JF, Cytrynbaum EN, Wasteneys GO. A CLASP-modulated cell edge barrier mechanism drives cell-wide cortical microtubule organization in Arabidopsis. Nat Commun. 2011 Aug 16;2(1):430.

9. Le PY, Ambrose C. CLASP promotes stable tethering of endoplasmic microtubules to the cell cortex to maintain cytoplasmic stability in Arabidopsis meristematic cells. Bassham DC, editor. PLoS ONE. 2018 Jun 12;13(6):e0198521.

10. Yu B, Zheng W, Xing L, Zhu JK, Persson S, Zhao Y. Root twisting drives halotropism via stress-induced microtubule reorientation. Developmental Cell. 2022 Oct;57(20):2412–2425.e6.

11. Dogterom M, Koenderink GH. Actin–microtubule crosstalk in cell biology. Nat Rev Mol Cell Biol. 2019 Jan;20(1):38–54.

12. Pimm ML, Henty-Ridilla JL. New twists in actin–microtubule interactions. Bement W, editor. MBoC. 2021 Feb 1;32(3):211–7.

13. Schlientz AJ, Bowerman B. C. elegans CLASP/CLS-2 negatively regulates membrane ingression throughout the oocyte cortex and is required for polar body extrusion. Colaiácovo MP, editor. PLoS Genet. 2020 Oct 7;16(10):e1008751.

14. Al-Bassam J, Chang F. Regulation of microtubule dynamics by TOG-domain proteins XMAP215/Dis1 and CLASP. Trends in Cell Biology. 2011 Oct;21(10):604– 14.

15. Akhmanova A, Steinmetz MO. Control of microtubule organization and dynamics: two ends in the limelight. Nat Rev Mol Cell Biol. 2015 Dec;16(12):711–26.

16. Macaisne N, Bellutti L, Laband K, Edwards F, Pitayu-Nugroho L, Gervais A, et al. Synergistic stabilization of microtubules by BUB-1, HCP-1, and CLS-2 controls microtubule pausing and meiotic spindle assembly. eLife. 2023 Feb 17;12:e82579.

17. Howe M, McDonald KL, Albertson DG, Meyer BJ. Him-10 Is Required for Kinetochore Structure and Function on Caenorhabditis elegans Holocentric Chromosomes. Journal of Cell Biology. 2001 Jun 11;153(6):1227–38.

18. Monen J, Maddox PS, Hyndman F, Oegema K, Desai A. Differential role of CENP-A in the segregation of holocentric C. elegans chromosomes during meiosis and mitosis. Nat Cell Biol. 2005 Dec;7(12):1248–55.

19. Dumont J, Oegema K, Desai A. A kinetochore-independent mechanism drives anaphase chromosome separation during acentrosomal meiosis. Nat Cell Biol. 2010 Sep;12(9):894–901.

20. Bel Borja L, Soubigou F, Taylor SJP, Fraguas Bringas C, Budrewicz J, Lara-Gonzalez P, et al. BUB-1 targets PP2A:B56 to regulate chromosome congression during meiosis I in C. elegans oocytes. eLife. 2020 Dec 23;9:e65307.

21. Dorn JF, Zhang L, Paradis V, Edoh-Bedi D, Jusu S, Maddox PS, et al. Actomyosin Tube Formation in Polar Body Cytokinesis Requires Anillin in C. elegans. Current Biology. 2010 Nov;20(22):2046–51.

22. Maddox AS, Azoury J, Dumont J. Polar body cytokinesis. Cytoskeleton. 2012 Nov;69(11):855–68.

23. Fabritius AS, Flynn JR, McNally FJ. Initial diameter of the polar body contractile ring is minimized by the centralspindlin complex. Developmental Biology. 2011 Nov;359(1):137–48.

24. Flynn JR, McNally FJ. A casein kinase 1 prevents expulsion of the oocyte meiotic spindle into a polar body by regulating cortical contractility. Chang F, editor. MBoC. 2017 Sep;28(18):2410–9.

25. Yan VT, Narayanan A, Wiegand T, Jülicher F, Grill SW. A condensate dynamic instability orchestrates actomyosin cortex activation. Nature. 2022 Sep 15;609(7927):597–604.

26. Ellefson, Marina L., McNally, Francis J. Kinesin-1 and Cytoplasmic Dynein Act Sequentially to Move the Meiotic Spindle to the OOcyte Cortex in Caenorhabditis elegans. Mol Biol Cell. 2009;20:2722–30.

27. Gigant E, Stefanutti M, Laband K, Gluszek-Kustusz A, Edwards F, Lacroix B, et al. Inhibition of ectopic microtubule assembly by the kinesin-13 KLP-7MCAK prevents chromosome segregation and cytokinesis defects in oocytes. Development. 2017 Jan 1;dev.147504.

28. Chuang CH, Schlientz AJ, Yang J, Bowerman B. Microtubule assembly and pole coalescence: early steps in *C. elegans* oocyte meiosis I spindle assembly. Biology Open. 2020 Jan 1;bio.052308.

29. 29. Harvey AM, Chuang CH, Sumiyoshi E, Bowerman B. C. elegans XMAP215/ZYG-9 and TACC/TAC-1 act at multiple times during oocyte meiotic spindle assembly and promote both spindle pole coalescence and stability. Colaiácovo MP, editor. PLoS Genet. 2023 Jan 6;19(1):e1010363.

30. Cheeseman IM, MacLeod I, Yates JR, Oegema K, Desai A. The CENP-F-like Proteins HCP-1 and HCP-2 Target CLASP to Kinetochores to Mediate Chromosome Segregation. Current Biology. 2005 Apr;15(8):771–7.

31. Maia ARR, Garcia Z, Kabeche L, Barisic M, Maffini S, Macedo-Ribeiro S, et al. Cdk1 and Plk1 mediate a CLASP2 phospho-switch that stabilizes kinetochore– microtubule attachments. Journal of Cell Biology. 2012 Oct 15;199(2):285–301.

32. Dumitru AMG, Rusin SF, Clark AEM, Kettenbach AN, Compton DA. Cyclin A/Cdk1 modulates Plk1 activity in prometaphase to regulate kinetochore-microtubule attachment stability. eLife. 2017 Nov 20;6:e29303.

33. Pereira C, Reis RM, Gama JB, Celestino R, Cheerambathur DK, Carvalho AX, et al. Self-Assembly of the RZZ Complex into Filaments Drives Kinetochore Expansion in the Absence of Microtubule Attachment. Current Biology. 2018 Nov;28(21):3408–3421.e8.

34. Laband K, Le Borgne R, Edwards F, Stefanutti M, Canman JC, Verbavatz JM, et al. Chromosome segregation occurs by microtubule pushing in oocytes. Nat Commun. 2017 Nov 14;8(1):1499.

35. Connolly AA, Sugioka K, Chuang CH, Lowry JB, Bowerman B. KLP-7 acts through the Ndc80 complex to limit pole number in C. elegans oocyte meiotic spindle assembly. Journal of Cell Biology. 2015 Sep 14;210(6):917–32.

36. Toyoda Y, Cattin CJ, Stewart MP, Poser I, Theis M, Kurzchalia TV, et al. Genome-scale single-cell mechanical phenotyping reveals disease-related genes involved in mitotic rounding. Nat Commun. 2017 Nov 2;8(1):1266.

37. Cartagena-Rivera AX, Logue JS, Waterman CM, Chadwick RS. Actomyosin Cortical Mechanical Properties in Nonadherent Cells Determined by Atomic Force Microscopy. Biophysical Journal. 2016 Jun;110(11):2528–39.

38. Zhovmer AS, Manning A, Smith C, Hayes JB, Burnette DT, Wang J, et al. Mechanical Counterbalance of Kinesin and Dynein Motors in a Microtubular Network Regulates Cell Mechanics, 3D Architecture, and Mechanosensing. ACS Nano. 2021 Nov 23;15(11):17528–48.

39. Farhadi L, Ricketts SN, Rust MJ, Das M, Robertson-Anderson RM, Ross JL. Actin and microtubule crosslinkers tune mobility and control co-localization in a composite cytoskeletal network. Soft Matter. 2020;16(31):7191–201.

40. Pintard L, Bowerman B. Mitotic Cell Division in *Caenorhabditis elegans*. Genetics. 2019 Jan 1;211(1):35–73.

41. O’Rourke SM, Christensen SN, Bowerman B. Caenorhabditis elegans EFA-6 limits microtubule growth at the cell cortex. Nat Cell Biol. 2010 Dec;12(12):1235–41.

42. Sugioka K, Bowerman B. Combinatorial Contact Cues Specify Cell Division Orientation by Directing Cortical Myosin Flows. Developmental Cell. 2018 Aug;46(3):257–270.e5.

43. Sulston JE, Schierenberg E, White JG, Thomson JN. The embryonic cell lineage of the nematode Caenorhabditis elegans. Developmental Biology. 1983 Nov;100(1):64–119.

44. Sepers JJ, Ramalho JJ, Kroll JR, Schmidt R, Boxem M. ERM-1 Phosphorylation and NRFL-1 Redundantly Control Lumen Formation in the C. elegans Intestine. Front Cell Dev Biol. 2022 Feb 7;10:769862.

45. Deng M, Li R. Sperm Chromatin-Induced Ectopic Polar Body Extrusion in Mouse Eggs after ICSI and Delayed Egg Activation. Bridger JM, editor. PLoS ONE. 2009 Sep 29;4(9):e7171.

46. Uraji J, Scheffler K, Schuh M. Functions of actin in mouse oocytes at a glance. Journal of Cell Science. 2018 Nov 15;131(22):jcs218099.

47. Harasimov K, Uraji J, Mönnich EU, Holubcová Z, Elder K, Blayney M, et al. Actin-driven chromosome clustering facilitates fast and complete chromosome capture in mammalian oocytes. Nat Cell Biol. 2023 Mar;25(3):439–52.

48. Olson SK, Greenan G, Desai A, Müller-Reichert T, Oegema K. Hierarchical assembly of the eggshell and permeability barrier in C. elegans. Journal of Cell Biology. 2012 Aug 20;198(4):731–48.

49. Johnston WL, Dennis JW. The eggshell in the C. elegans oocyte-to-embryo transition. Genesis. 2012 Apr;50(4):333–49.

50. Zhang X, Ma C, Miller AL, Katbi HA, Bement WM, Liu XJ. Polar Body Emission Requires a RhoA Contractile Ring and Cdc42-Mediated Membrane Protrusion. Developmental Cell. 2008 Sep;15(3):386–400.

51. Deng M, Suraneni P, Schultz RM, Li R. The Ran GTPase Mediates Chromatin Signaling to Control Cortical Polarity during Polar Body Extrusion in Mouse Oocytes. Developmental Cell. 2007 Feb;12(2):301–8.

52. Hubbard EJA, Greenstein D. TheCaenorhabditis elegans gonad: A test tube for cell and developmental biology. Dev Dyn. 2000 May;218(1):2–22.

53. Bembenek JN, Richie CT, Squirrell JM, Campbell JM, Eliceiri KW, Poteryaev D, et al. Cortical granule exocytosis in *C. elegans* is regulated by cell cycle components including separase. Development. 2007 Nov 1;134(21):3837–48.

54. Gilliland WD, Hughes SE, Cotitta JL, Takeo S, Xiang Y, Hawley RS. The Multiple Roles of Mps1 in Drosophila Female Meiosis. Barsh G, editor. PLoS Genet. 2007 Jul 13;3(7):e113.

55. Bender M, Thon JN, Ehrlicher AJ, Wu S, Mazutis L, Deschmann E, et al. Microtubule sliding drives proplatelet elongation and is dependent on cytoplasmic dynein. Blood. 2015 Jan 29;125(5):860–8.

56. Patel-Hett S, Richardson JL, Schulze H, Drabek K, Isaac NA, Hoffmeister K, et al. Visualization of microtubule growth in living platelets reveals a dynamic marginal band with multiple microtubules. Blood. 2008 May 1;111(9):4605–16.

57. Kirik V, Herrmann U, Parupalli C, Sedbrook JC, Ehrhardt DW, Hülskamp M. CLASP localizes in two discrete patterns on cortical microtubules and is required for cell morphogenesis and cell division in *Arabidopsis*. Journal of Cell Science. 2007 Dec 15;120(24):4416–25.

58. Meiring JCM, Grigoriev I, Nijenhuis W, Kapitein LC, Akhmanova A. Opto-katanin, an optogenetic tool for localized, microtubule disassembly. Current Biology. 2022 Nov;32(21):4660–4674.e6.

59. Li Y, Munro E. Filament-guided filament assembly provides structural memory of filament alignment during cytokinesis. Developmental Cell. 2021 Sep;56(17):2486–2500.e6.

