## Supplementary figures and images for "Cortical microtubules oppose actomyosin-driven membrane ingression during *C. elegans* meiosis I polar body extrusion"

### Supplemental 1

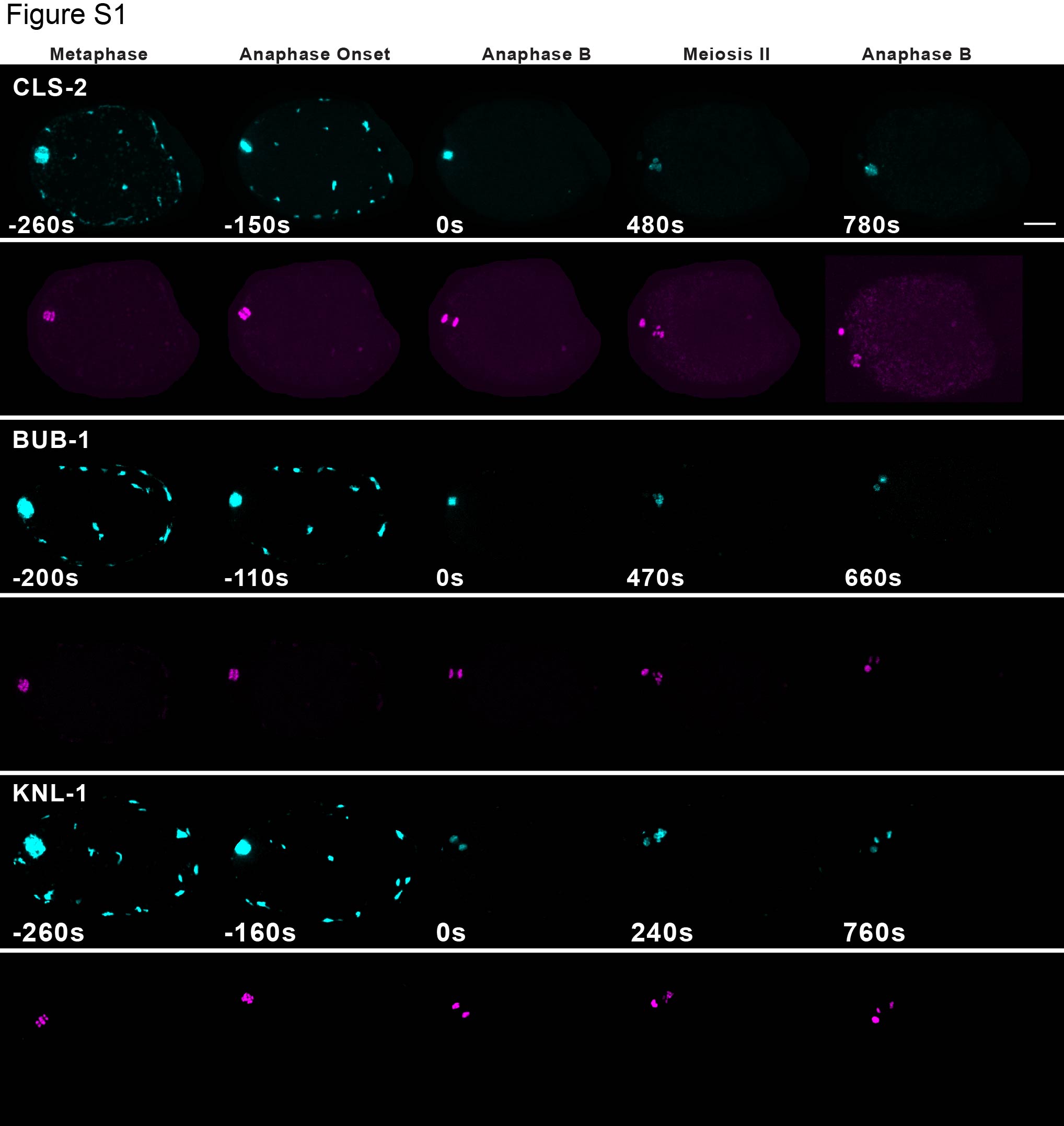

### Supplemental 2

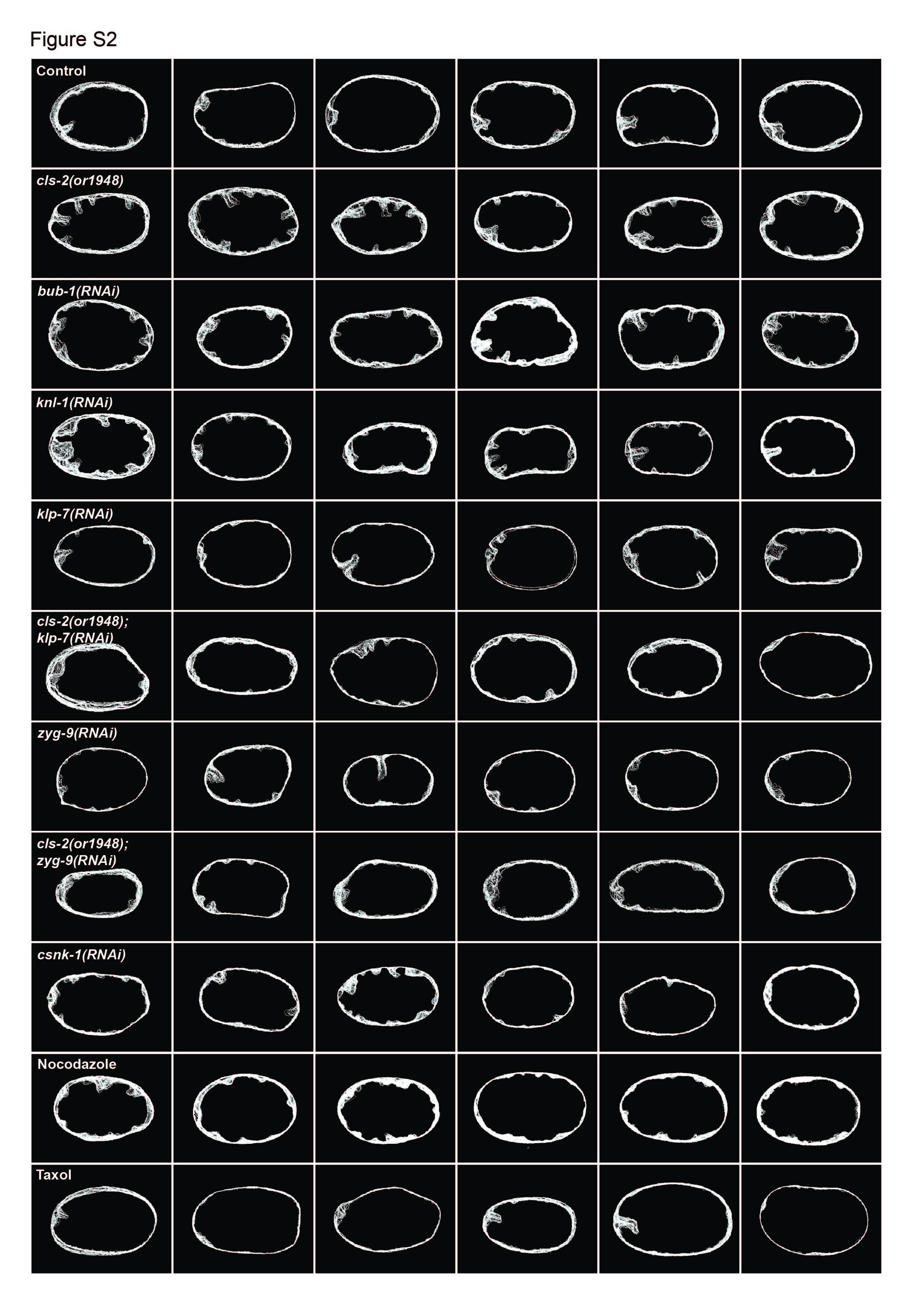

### Supplemental 3

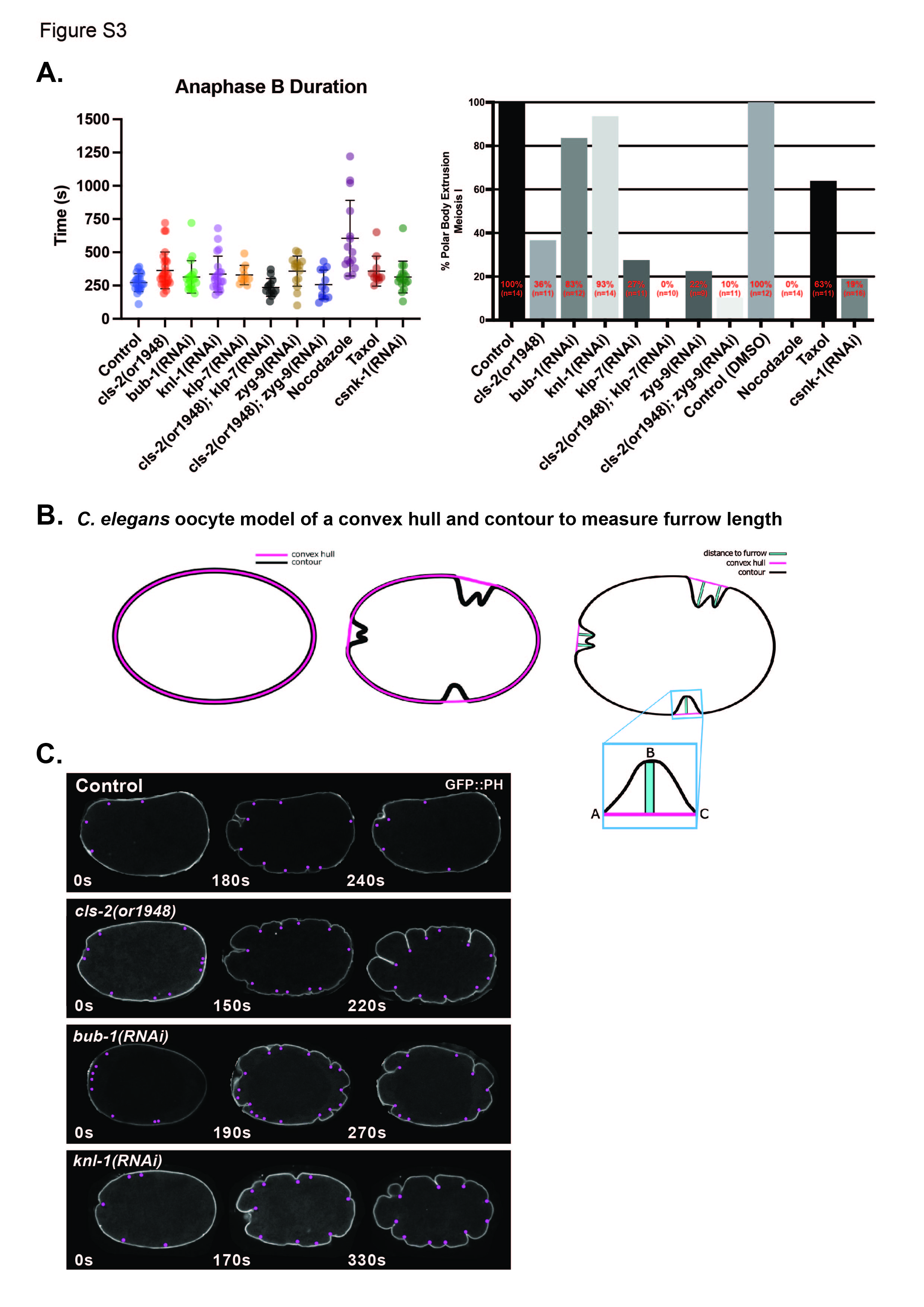

### Supplemental 4

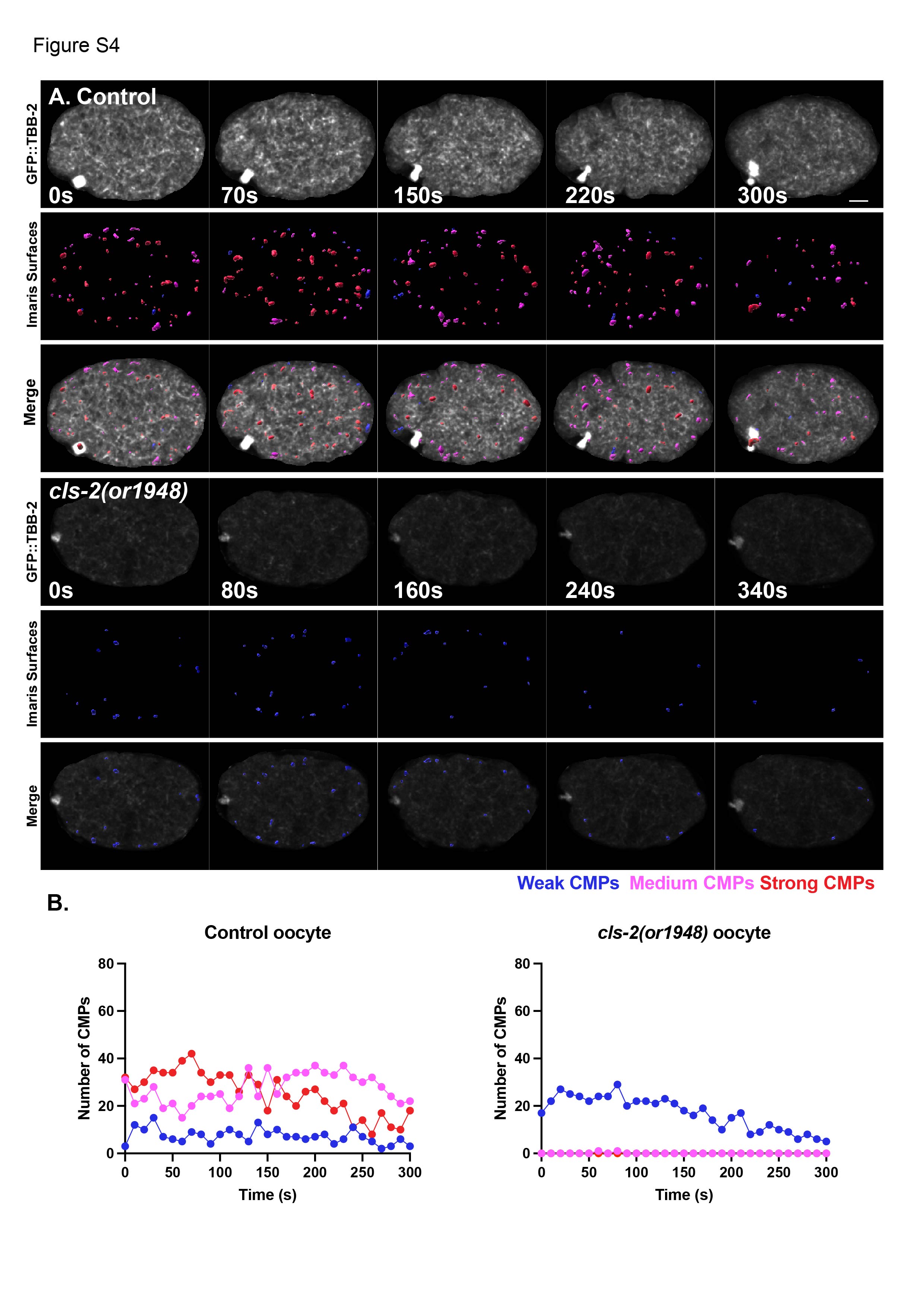

### Supplemental 5

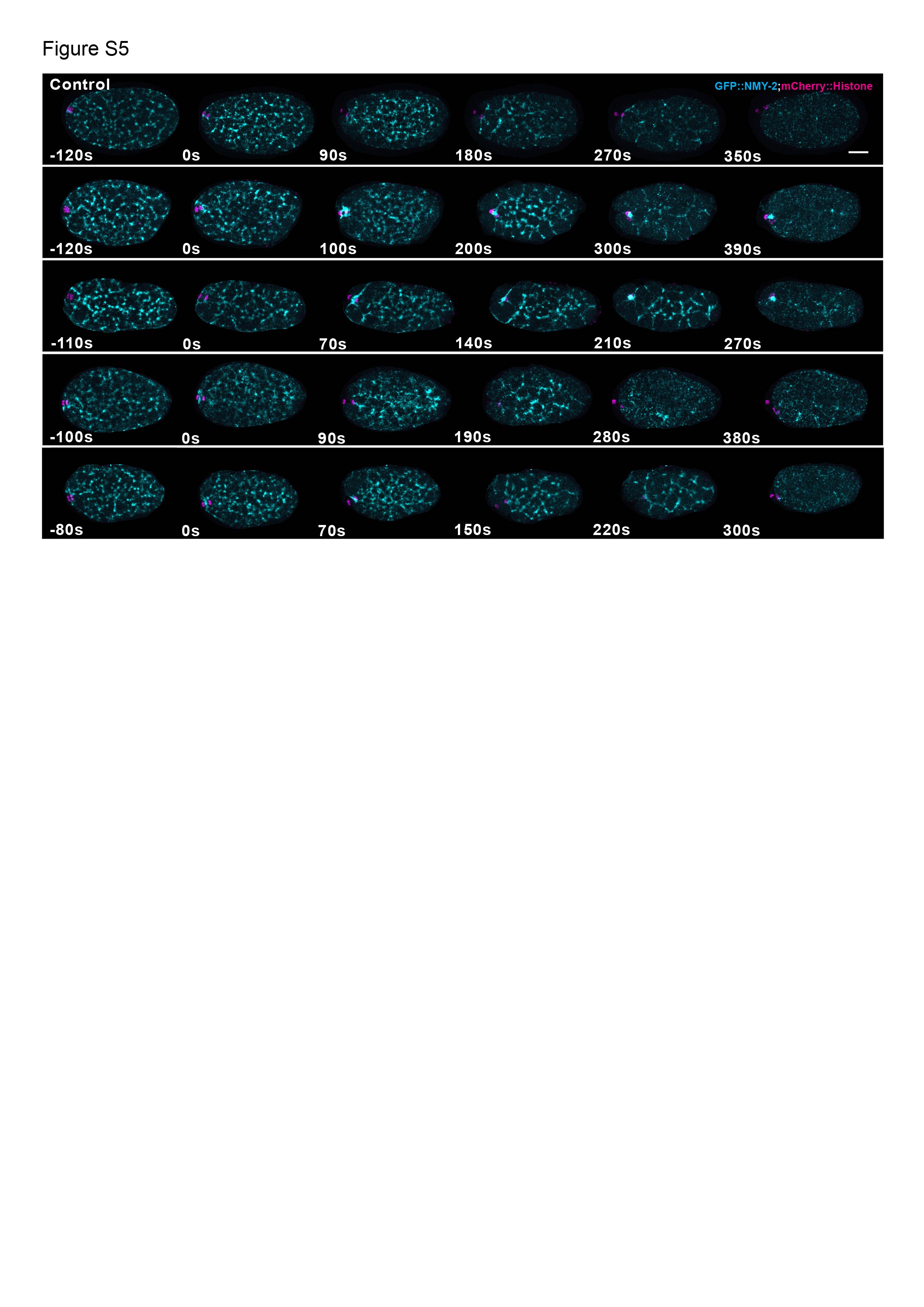

### Supplemental 6

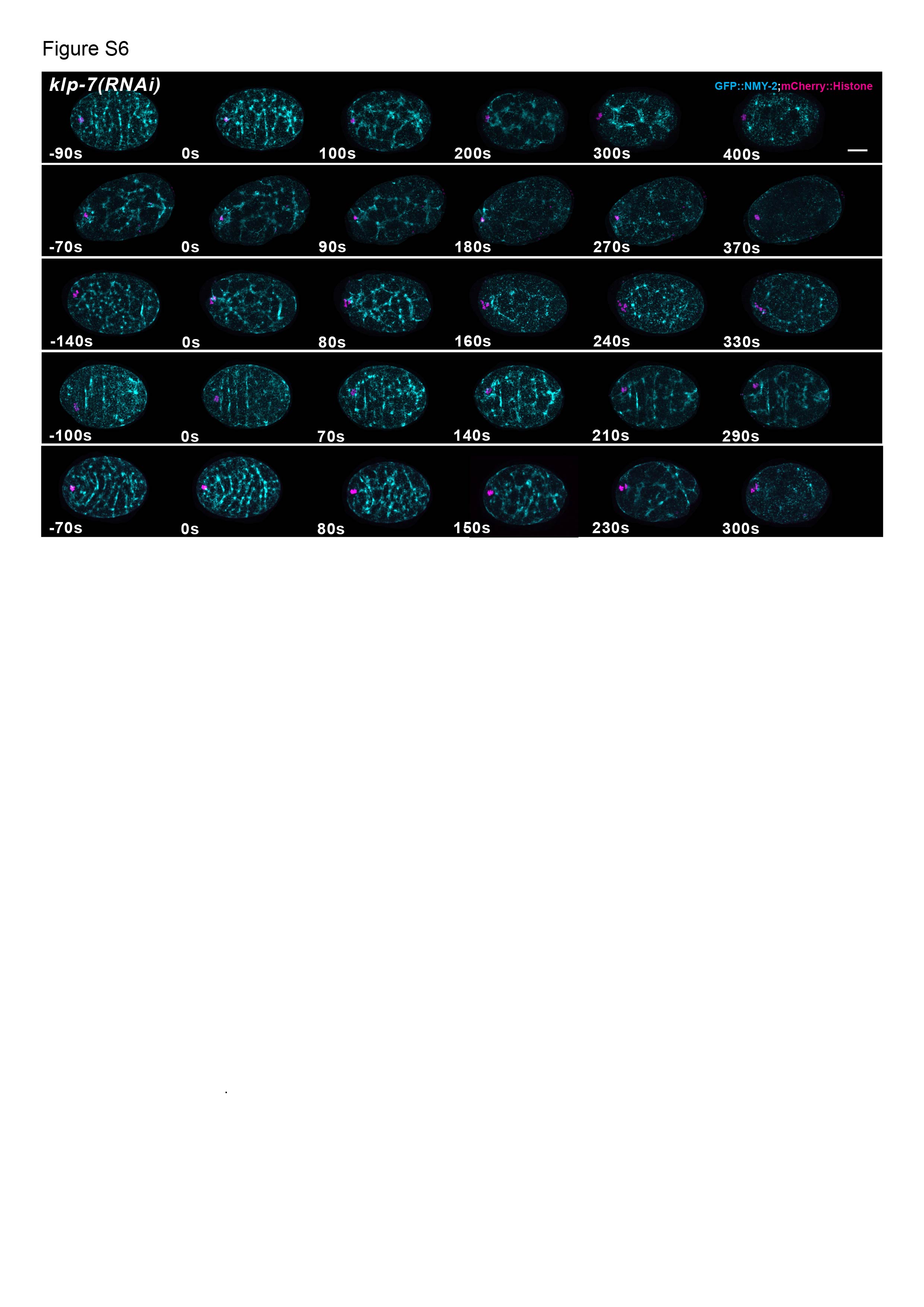

### Supplemental 7

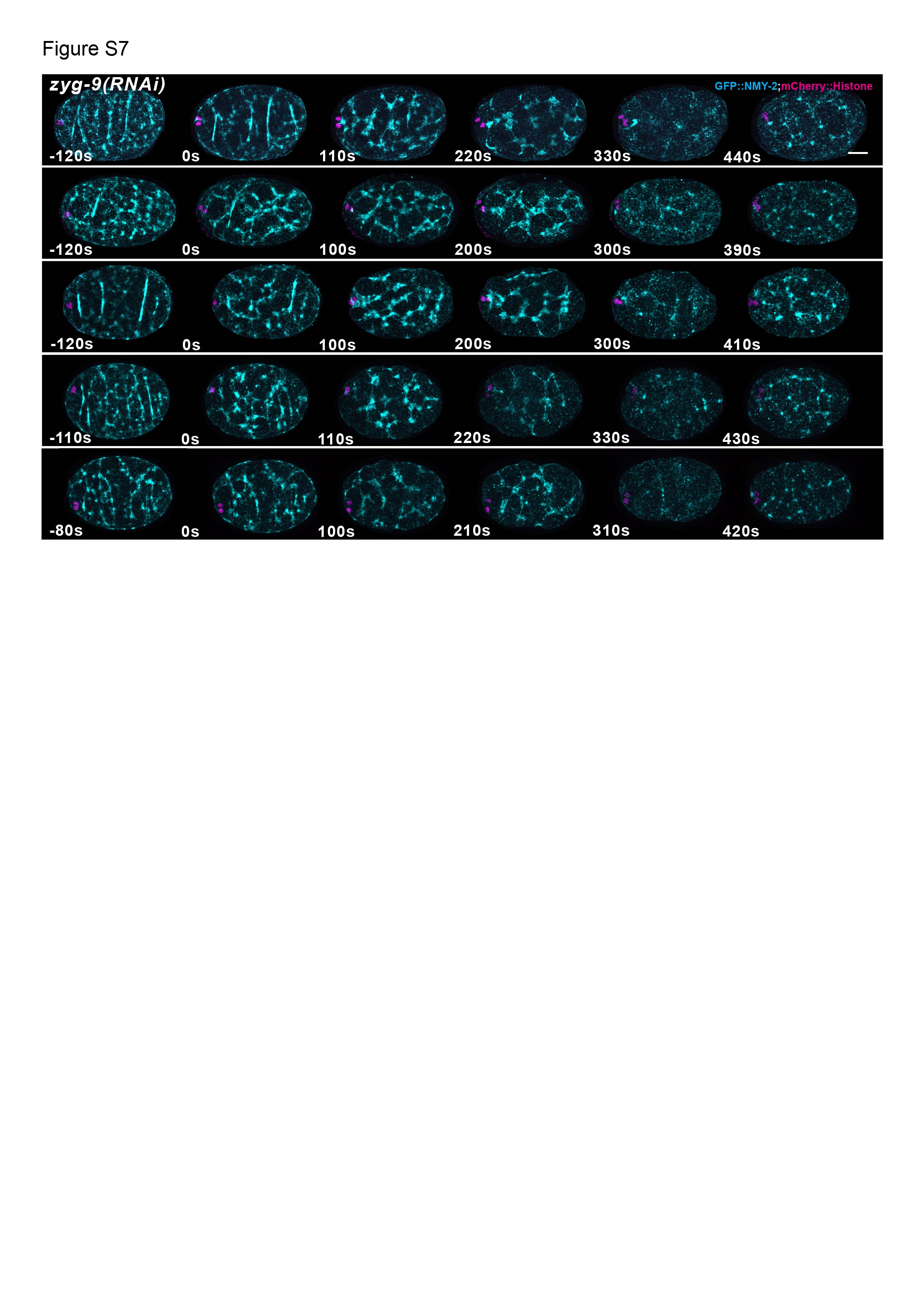

### Supplemental 8

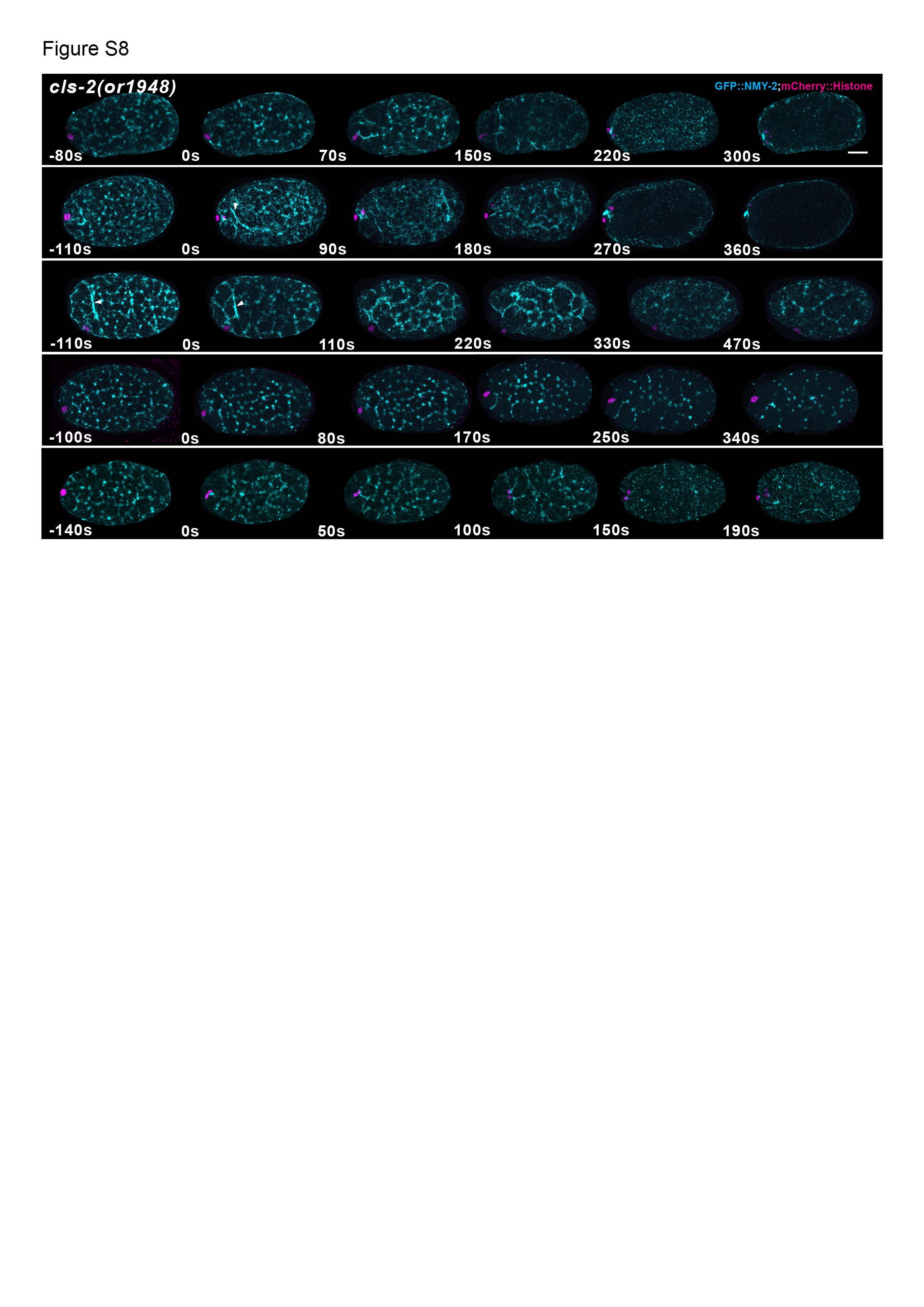

### Supplemental 9

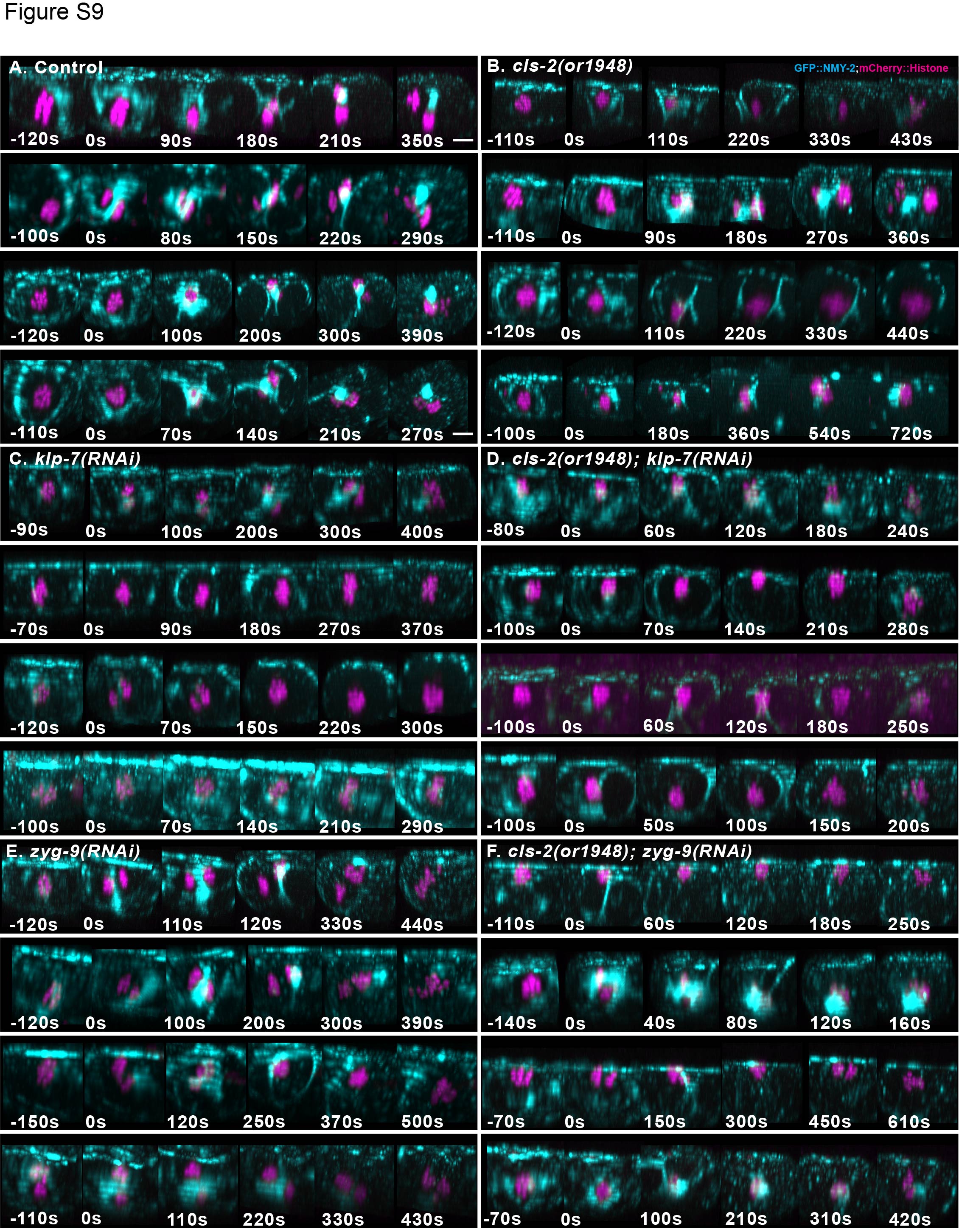

### Supplemental 10

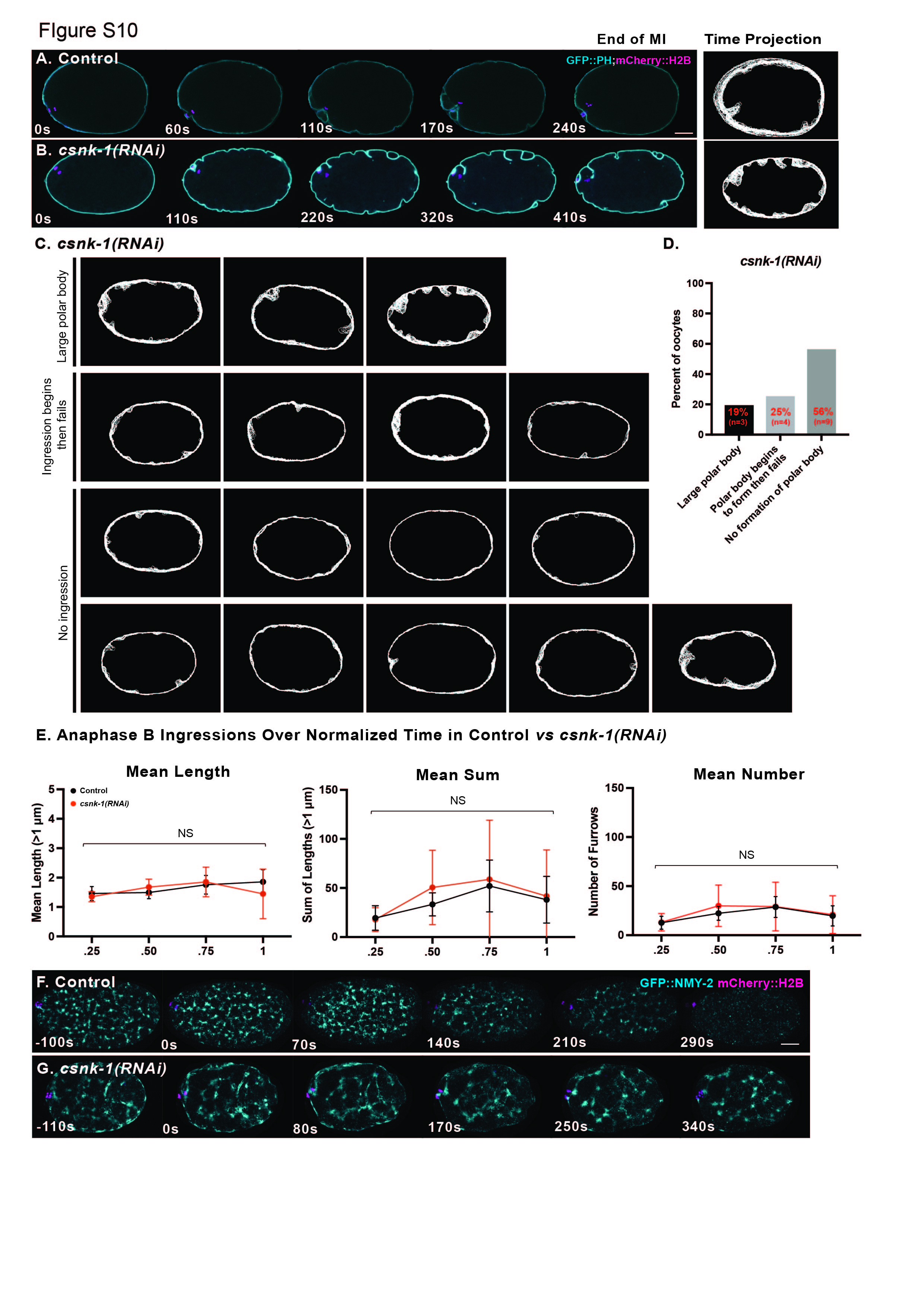
